# LSD1 promotes secretory cell specification to drive *BRAF* mutant colorectal cancer

**DOI:** 10.1101/2020.09.25.313536

**Authors:** Samuel A. Miller, Robert A. Policastro, Shruthi Sriramkumar, Tim Lai, Thomas D. Huntington, Christopher A Ladaika, Gabriel E. Zentner, Heather M. O’Hagan

**Affiliations:** Genome, Cell, and Developmental Biology, Department of Biology, Indiana University Bloomington, Bloomington, IN, 47405, USA; Medical Sciences Program, Indiana University School of Medicine, Bloomington, IN, 47405, USA; Cell, Molecular and Cancer Biology Graduate Program, Indiana University School of Medicine, Bloomington, IN, 47405, USA; Luddy School of Informatics, Computing, and Engineering, Indiana University, Bloomington, IN, 7408, USA; Department of Mathematics, Indiana University, Bloomington, IN, 47405, USA; Indiana University Melvin and Bren Simon Comprehensive Cancer Center, Indianapolis, IN, 46202, USA

## Abstract

Despite the connection to distinct mucus-containing colorectal cancer (CRC) histological subtypes, the role of secretory cells, including goblet and enteroendocrine (EEC) cells, in CRC progression has been underexplored. Analysis of TCGA and single cell RNA sequencing data demonstrates that multiple secretory progenitor populations are enriched in *BRAF-*mutant CRC patient tumors and cell lines. Enrichment of EEC progenitors in *BRAF*-mutant CRC is maintained by DNA methylation and silencing of *NEUROD1*, a key gene required for differentiation of EECs. Mechanistically, secretory cells and the factors they secrete, such as Trefoil factor 3, are shown to promote colony formation and activation of cell survival pathways in the entire cell population. We further identify LSD1 as a critical regulator of secretory cell specification *in vitro* and in a colon orthotopic xenograft model, where LSD1 loss reduces tumor growth and metastasis. This work establishes EEC progenitors, in addition to goblet cells, as targetable populations in *BRAF*-mutant CRC and identifies LSD1 as a therapeutic target in secretory lineage-containing CRC.

## Introduction

The human large intestine is populated by the stem and transit-amplifying cells that give rise to differentiated absorptive enterocytes and secretory cells such as goblet and enteroendocrine (EEC) cells. Renewal of these populations in the adult intestine relies on a delicate balance between proliferation and differentiation. This equilibrium is disrupted in colorectal cancer (CRC), canonically marked by highly proliferative and moderately differentiated cells(Fleming et al., 2012). CRC that exhibits mucin hypersecretion that encompasses greater than 50% of the total tumor volume is classified as mucinous adenocarcinomas(Bosman et al., 2010). Approximately 15% of CRC falls into this pathologic classification(Hugen et al., 2014), indicating that a sizable fraction of CRC patients exhibit some degree of secretory cell differentiation and suggesting that secretory cells may be important in driving a proportion of CRC. However, because these pathological diagnoses are based on levels of secreted mucin, they may not reflect the true proportion or diversity of secretory cells in CRC. Tumors classified as non-mucinous may still contain secretory cells and/or all tumors classified as mucinous may not contain the same subset of secretory cells. Substantial heterogeneity in these groups could potentially explain previously noted conflicting results regarding the survival of patients with mucinous versus non-mucinous CRC(Schneider and Langner, 2014).

Another common subtype of cancer originating from secretory cells are neuroendocrine tumors (NETs), which arise from endocrine lineage cells. NETs are less common in the large intestine but are the most prevalent cancer subtype of the appendix (Moertel et al., 1968) (Modlin and Sandor, 1997). Aggressive subtypes of NETs include those with goblet cell differentiation (goblet cell carcinoids)(McGory et al., 2005), and poorly differentiated NETs (PDNETs)(Basturk et al., 2014). While distinct from mucinous tumors, goblet cell carcinoids are pathologically more similar to adenocarcinomas of the large intestine than to other NET subtypes(van Eeden et al., 2007), which may contribute to their under-characterization in the large intestine due to the difficulty in differentiating them from canonical adenocarcinomas. However, recent pathological studies have begun to identify tumors similar to goblet cell carcinoids in the large intestine(Inoue et al., 2020) (Abdalla et al., 2020), indicating that mixed secretory lineage CRCs form at some previously unappreciated frequency. It is currently unknown whether goblet and EEC cells present in colon adenocarcinomas functionally contribute to tumor progression. In spite of this knowledge gap, mounting evidence suggests that oncogenic transformation may be followed by altered differentiation of secretory lineages(Peignon et al., 2011).

Driver mutations in CRC categorically affect essential cell processes, such as cell fate, cell survival, and/or genome maintenance(Vogelstein et al., 2013). Loss of function mutations in the *Adenomatous Polyposis Coli* gene (*APC*) induce non-canonical β-catenin-mediated degradation of the general secretory specification factor Atonal Homolog 1 (ATOH1) leading to loss of goblet and presumably EEC lineage cells(Peignon et al., 2011). Alternatively, mucinous tumors have increased mutation rates in genes that function in the RAS/MAPK and PI3K/AKT signaling pathways(Schneider and Langner, 2014). Accordingly, inducing the *BRAF(V600E)* mutation in mouse colon organoids increased mucin biosynthesis(Reischmann et al., 2020), but the mechanism by which *BRAF* mutation alters the activity or differentiation of secretory cells is poorly understood.

The chromatin-modifying enzyme lysine-specific demethylase 1 (*LSD1*) is over-expressed in many CRCs and contributes to the formation of numerous cancer types such as small cell lung cancer and acute myeloid leukemia(Mohammad et al., 2015) (Takagi et al., 2017) (Maiques-Diaz et al., 2018). While the demethylase activity of LSD1 contributes to the progression of multiple cancer types, the scaffolding function of LSD1 in the CoREST complex has recently been highlighted in CRC(Miller et al., 2020). Nevertheless, both the normal functions of LSD1 in the intestine and its potential role in intestinal cancers are poorly understood. LSD1 has been directly implicated as a regulator of neuronal and myeloid differentiation, brown adiposities, and muscle cells(Laurent et al., 2015) (Maiques-Diaz et al., 2018) (Lin and Farmer, 2016) (Tosic et al., 2018). Furthermore, recent studies have begun to connect LSD1 with the regulation of intestinal secretory lineages. Co-transfection of *LSD1* with the EEC lineage transcription factor *NEUROD1* upregulated expression of the EEC marker *SCT* in HuTu80 and HEK 293 cells, indicating that LSD1 may function in the regulation of EEC lineage genes(Ray et al., 2014). Another study found that while LSD1 was required for Paneth cell formation during mouse development, loss of LSD1 did not significantly affect goblet or EEC formation in the small intestine(Zwiggelaar et al., 2020). An outstanding question in the field is whether the well-established tumor-promoting function of LSD1 in the intestine may be linked to an undescribed role in intestinal cell differentiation.

In this study, using patient data and *in vitro* culture systems, we discover that a subset of *BRAF* kinase mutated CRC are enriched for EECs that are locked in the progenitor state due to a DNA methylation-dependent differentiation blockade. We generate single-cell RNA-sequencing (scRNA-seq) datasets to study the regulation and function of EEC progenitors in CRC as well as in comparison to the normal human colon at high resolution, identifying conserved and CRC specific features. We further determine that LSD1 is essential for the formation of both EEC and goblet cell lineages in *BRAF* tumors and that loss of these populations abrogates tumor growth and metastases *in vivo*. Overall, this study establishes EEC progenitors, in addition to goblet cells, as targetable populations in *BRAF*-mutant CRC and identifies LSD1 as a novel therapeutic avenue to deplete EEC and goblet lineage cells, commonly enriched in multiple subtypes of *BRAF* mutant CRC.

## Methods

### Data availability

Single-cell RNA-seq data generated in this study as well as Wang *et al.,* 2019 can be explored at http://185.215.224.30:3838/miller/miller_policastro_2020

(*permanent address pending*). Raw scRNA-seq datasets are available through NCBI’s Gene Expression Omnibus (GEO) through GEO series accession number (*pending, available on request*). Processed scRNA-seq datasets can be accessed through the BROAD Single cell portal. We developed an R library to calculate the proportional difference in cell number proportions between clusters in scRNA-seq, available at https://github.com/rpolicastro/scProportionTest/releases/tag/v1.0.0. Bulk RNA-sequencing data in HT29 and SW480 cells were accessed using NCBI’s Gene Expression Omnibus (GEO) through GEO series accession number GSE139927.

### Single-cell sequencing and TCGA analyses

10,000 cells per sample were targeted for input to the 10X Genomics Chromium system using the Chromium Next GEM Single Cell 3′ Kit v3.1 at the IU School of Medicine (IUSM) Center for Medical Genomics core (HT29) or at IU Bloomington (H508). The libraries were sequenced at the IUSM Center for Medical Genomics using a NovaSeq 6000 with a NovaSeq S2 reagent kit v1.0 (100 cycles) with approximately 450 million read pairs per sample. A detailed description of single-cell sequencing analysis as well as TCGA analysis is included in the supplemental methods.

### Cell culture and treatments

All cell lines were maintained in a humidified atmosphere with 5% CO_2_. HT29 cells were cultured in McCoys 5A media (Corning), NCI-H508 in RPMI 1640 media (Corning) and LS-174T cells in DMEM, supplemented with 10% FBS (Gibco). All cell lines were purchased from the ATCC; LS174T (2020), NCI-H508 (2019), HT29 (verified by STR profiling in 2019). Decitabine (DAC) (Sigma A3656) was reconstituted in sterile ddH_2_O at 2mg/ml, and administered at 100nM in fresh media every 24 hours for a total of 72 hours. ISX-9 (TOCRIS, 4439) was reconstituted in DMSO at 50mM, and cells were treated with 40μM for 24 hours. For DAC/ISX-9 co-treatment, cells were treated with 100nM DAC or ddH_2_O for 48 hours, and on day three, treated with 100nm DAC or ddH_2_0 and 40μM ISX-9 or DMSO for 24 hours. For Notch inhibition experiments, cells were treated once with 40nM DBZ solubilized in DMSO (Tocris, 4489) for 48H. For TFF3 and EGF combination treatments in media exchange experiments, cells were grown for 48H in serum-free media, which was then collected and filtered with a .45 micron filter. 1μg/ml anti-TFF3 (R&D Systems, MAB4407) or IgG was added to the media for 5 minutes. These compounds were solubilized in PBS prior to treatment. The media was then added back to the EV or LSD1 KD cells for 24 hours. Cells were then treated with 125 ng/ml recombinant EGF (R&D Systems: 236-EG) for 1 hour and collected for western blot analysis. For CellTiter-Glo® assay (Promega #G7572) involving combination treatment, cells were starved in media lacking serum for 48H prior to treatment. Cells were treated with 1 μg/ml anti-TFF3 or IgG control and 125 ng/ml recombinant EGF for 48H in serum-free media.

### TFF3 ELISA and MTT assay

For TFF3 ELISA, cells were cultured in serum-free media for 24H. Fresh serum-free media was added, and after an additional 24H, was collected and centrifuged at 16,000xg for 5 minutes at 4°C. ELISA’s were performed using 50 μl of media per sample and following the manufacturer’s protocol (R&D systems, DTFF30). After removal of media, 1 ml of Optimem + 100 μl MTT (5 mg/ml in PBS) was added and plates were incubated at 37°C for 3.5H. The media was removed and DMSO was added. Absorbance was read at 570nM.

### RNA isolation for qPCR

Total RNA was isolated using the RNeasy mini kit (Qiagen, 74104). For qPCR, RNA was used to generate cDNA via reverse transcription (Thermo, K1642). cDNA was amplified using gene-specific primers and FastStart Essential DNA Green Master (Roche, 06402712001). Cq values of non-housekeeping genes were normalized to *RHOA or b-ACTIN* expression, selected using the Housekeeping Transcript Atlas (housekeeping.unicamp.br) as strong colon housekeeping genes. qPCR Primer sequences are included as supplemental materials. Results are shown as mean +/− SD.

### Clonogenic growth and proliferation assays

For clonogenic growth assays, 500 single cells were plated on one well of a 6-well plate and allowed to grow for 14 days at 37 °C. Cells were then fixed with 10% formalin and stained with crystal violet. Stained cells were imaged using an SYNGENE G:BOX and quantification were carried out using the GeneSys and GeneTools programs. Proliferation assays were performed using the CellTiter-Glo® Luminescent Cell Viability Assay (Promega #G7572) per manufacturer’s protocol. Briefly, 1×10^3^ cells were plated in 96 well plates and allowed to incubate under standard growth conditions. Results are shown as mean +/− SD.

### Knockdown

LSD1 (KDM1A) (TRCN0000327856, TRCN0000327932, TRCN0000046071), ATOH1 (MATH1) (TRCN0000433089, TRCN0000414973), NEUROG3 (TRCN0000427521) and SPDEF(TRCN0000274422) knockdown constructs were purchased from mission shRNA; empty plasmid was used as a vector control. Knockdowns were performed as previously described(Miller et al., 2020).

### Immunofluorescence and imaging

For tissue culture, cells were grown on coverslips at 37°C. Cells were fixed with 4% paraformaldehyde in PBS 15 minutes at room temperature (RT), then permeabilized with 0.5% Triton-X in PBS for 10 minutes. After permeabilization, cells were blocked with 1% BSA in PBST (PBS+ 0.2% Tween 20) for 30 minutes. Cells were then incubated with anti-LSD1 (CST, #2139,1:100), anti-TFF3 (R& D systems, MAB4407,1:100), anti-SUSD2 (Biolegend, #327406,1:100), anti-MUC2 (Fisher, PA5-21329,1:100), or anti-INSM1 (Santa Cruz, #271408,1:100) in 1% BSA in PBST for 1 hour at RT. This was followed by incubation with secondary antibody using Alexa Fluor conjugate (CST, rabbit, #4412, 1:1000, #8889,1:500) for 1 hour at RT. Coverslips were secured by Prolong Gold Antifade with DAPI (Cell Signaling Technology, #8961). Images were acquired on a Leica SP8 scanning confocal system with DMi8-inverted microscope with LASX software (Leica Microsystems). All the images were taken at either 40x or 63x magnification, 1.4A oil immersion at RT and processed using ImageJ (NIH).

### Whole-cell isolation and western blot

For protein isolation, cell pellets were lysed in a 4% SDS buffer using a Qiashredder (Qiagen). Relative densitometry for western blots was determined using ImageJ software and normalized to the density of loading control β-ACTIN. Results are shown as mean +/− SD.

### Quantitative Methylation Specific PCR (qMSP)

Total DNA was isolated using the DNeasy kit (Qiagen 69504). DNA was bisulfite-treated (EZ DNA methylation lightning kit, Zymo Research, D5030) and used for qMSP. qMSP assays were first validated using a standard curve of bisulfite-treated mixtures of unmethylated and methylated DNA (data not shown). To calculate relative DNA methylation, qMSP assays were performed for each gene using primer sets specific for unmethylated (U) and methylated (M) alleles. The delta Cq method was used to calculate M/U.

### Orthotopic xenograft mouse model

HT29 cells were infected with the pLenti-U6-tdTomato-P2A-BlasR (LRT2B) virus (Addgene, 1108545) that enables dual production of firefly luciferase and dTomato. A strong dTomato signal was detectable within 14 days. Instead of sorting individual clones, the top 20% brightest dTomato cells were isolated by FACs to maintain the natural heterogeneity of these cells. 1,000,000 LSD1 KD or vector control cells were combined with Matrigel and injected into the mechanically prolapsed mouse large intestine. To visualize the firefly luciferase positive cells *in vivo*, mice were injected with luciferin and imaged using an IVIS imaging system every 7 days. After approximately five weeks, mice were sacrificed and xenograft primary tumors along with adjacent normal mouse tissue, lungs and livers were harvested, imaged using an IVIS imaging system, fixed in 4% paraformaldehyde, and frozen in OTC for further analysis. For TFF3 immunofluorescence, tissue sections were incubated with anti-TFF3 (R&D Systems, MAB4407, 1:50) overnight at 4°C followed by Alexa Flour anti-mouse 488 (Cell Signaling Technology, 4412, 1:500) for 1 hour at RT. For alcian blue staining, tissue sections were incubated in 3% acetic acid for 3 minutes, stained with Alcian Blue Solution (Electron Microscopy Sciences, 26323-01) for 30 minutes and counterstained with Nuclear Fast Red Solution (Electron Microscopy Sciences, 16323-02) for 5 minutes. All mouse experiments were covered under a protocol approved by the Indiana University Bloomington Animal Care and Use Committee in accordance with the Association for Assessment and Accreditation of Laboratory Animal Care International.

## Results

### BRAF mutation is associated with poorly differentiated EECs

While it is generally understood how differentiation is coordinated in the intestine, whether differentiation is altered in CRC is currently understudied. In canonical intestinal epithelium differentiation, absorptive progenitors are specified by HES1(Jensen et al., 2000), and cells without NOTCH1-mediated repression of *ATOH1* form the various secretory cells(Yang et al., 2001) (Figure 1A). ATOH1+ cells become goblet cells and NEUROG3+ cells, which become multipotent progenitors and eventually mature EECs(Schonhoff et al., 2004). The well-defined progenitor state is potentially unique to the EEC lineage, as goblet cells currently have no resolved progenitor stage. It is unknown, however, whether oncogenic mutations alter the specification of the EEC lineage in CRC.

**Figure 1:**
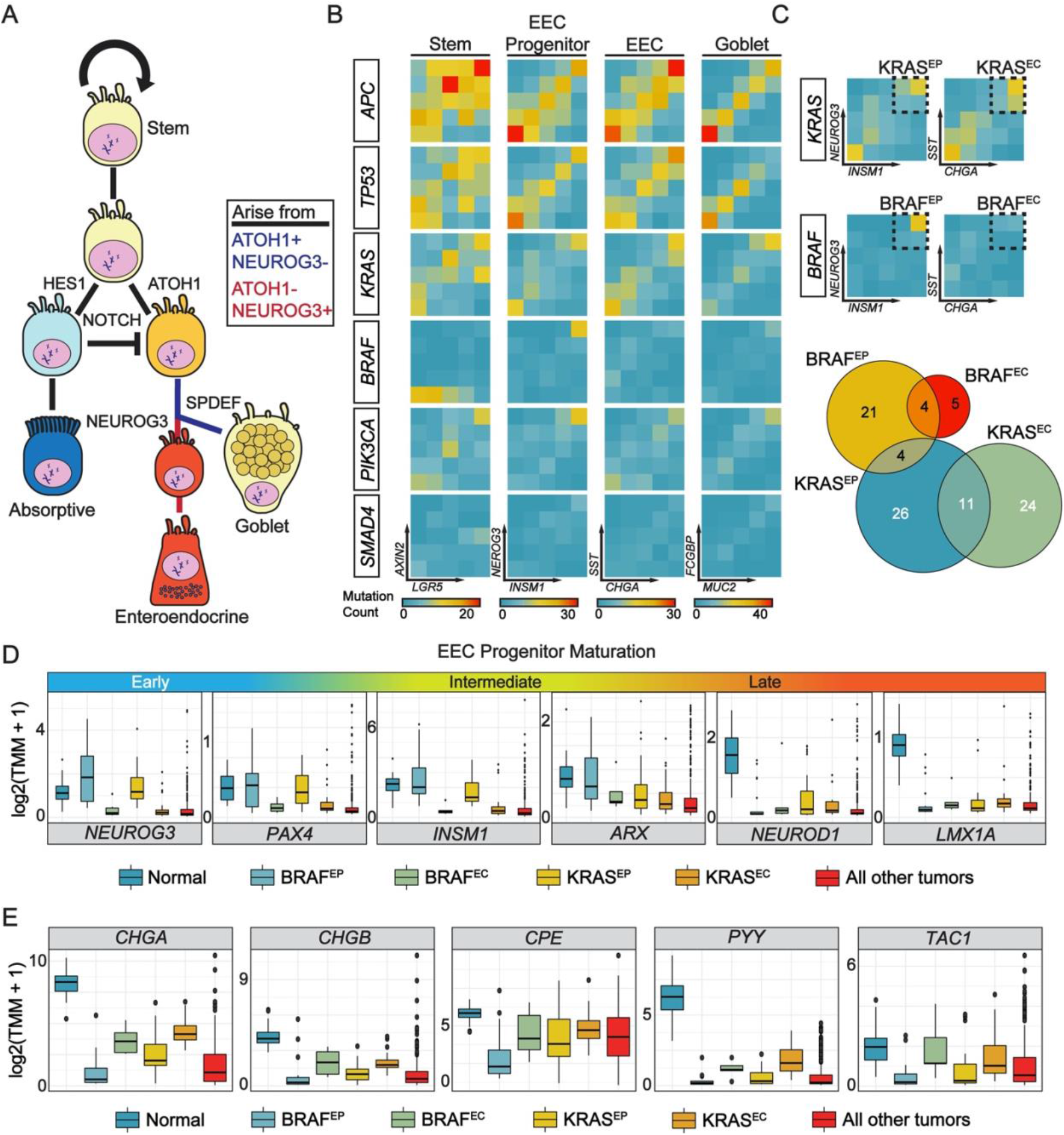
Enteroendocrine progenitors are enriched in a subset of *BRAF* mutant CRC. A, A diagrammatic representation of intestinal epithelium differentiation. B, Co-expression of cell-type marker genes plotted by driver mutation type and visualized through the overlap of normalized Log_2_ trimmed mean of M values (TMM) count quantiles. C, Diagram depicting the selection of BRAF^EP^, BRAF^EC^, KRAS^EP^, and KRAS^EC^ groups, as well as Euler plot of the number of samples for each group. D & E, Boxplot of gene expression in normal tissue (Normal), BRAF^EP^, BRAF^EC^, KRAS^EP^, KRAS^EC^, and all other COAD dataset tumors (“All other tumors”). Boxplot includes the median and box edges represent 2nd and 3rd quartiles and whiskers represent 1st and 4th quartiles.

To determine if the frequently mutated-in-CRC genes *KRAS*, *APC*, *TP53*, *BRAF*, *PIK3CA*, or *SMAD4* are associated with the differentiation status of secretory cells in CRC, we used expression and mutation data from the TCGA COAD (supplemental methods). Canonical marker genes *LGR5/AXIN2* (Stem), *INSM1/NEUROG3* (Enteroendocrine progenitor), *SST/CHGA* (Enteroendocrine), and *MUC2/FCGBP* (Goblet) were all significantly correlated (p-value < 0.05 and R > 0.45) (Supplemental Figure 1A). Thus, the coexpression of these marker gene pairs was used as an approximation of cell type content in the tumors. We annotated the expression quintiles of these marker pairs among all samples, and then plotted the samples by *KRAS*, *APC*, *TP53*, *BRAF*, *PIK3CA*, or *SMAD4* mutation (Figure 1B). For each mutation, we then computed the number of samples overlapping on each pairwise gene expression quintile and calculated FDR corrected p-values using a Monte Carlo simulation with 10,000 iterations (Supplemental Figure 1B).

Some trends of note include that *APC* mutations were most frequent and enriched in tumors where stem cell marker expression was high as well as in tumors where EEC progenitor or goblet cell marker expression was low (Figure 1B). Conversely, *BRAF* mutant tumors were enriched in tumors where stem cell marker expression was low. *BRAF* mutations were enriched in tumors with high expression of goblet cell markers, consistent with previous studies of mucinous CRC(Schneider and Langner, 2014). *BRAF* mutations were also highly enriched in tumors where EEC progenitor marker expression was exclusively high, a previously uncharacterized association. The clear positive association between *BRAF* mutation and the enrichment of EEC progenitor and goblet cells was not observed in other secretory lineage cell-types, such as Paneth or tuft cells (Supplemental Figure 1C-E). Furthermore, while absolute counts were similar between *KRAS*, *APC*, *TP53*, *PIK3CA*, and *SMAD4* mutants with high EEC progenitor and differentiated EEC marker expression, this was not the case with *BRAF* mutations, where high absolute counts and enrichment were only observed in progenitors. While *BRAF* and *KRAS* both function in the p38-RAS pathway, this analysis suggests that tumors harboring mutations in either gene exhibit different EEC cell differentiation statuses in CRC.

To study this apparent dichotomy, tumors were split into four groups bearing the *BRAF* or *KRAS* mutation, and expression values in the top two quintiles of mature EEC progenitor markers (BRAF^EP^/KRAS^EP^) or of differentiated EEC markers (BRAF^EC^/KRAS^EC^), (Figure 1C). *KRAS* mutation is equally common among tumors with high expression of progenitor or differentiated markers, but *BRAF* mutation is primarily associated with high expression of progenitor markers. Both *BRAF* and *KRAS* groups generally exhibited elevated levels of the secretory specification factor *ATOH1* and goblet marker *MUC2* compared to the all other tumors group (Supplemental Figure 1F & Supplemental Table 3). However, BRAF and KRAS groups had a similar distribution of pathological classifications to the all other tumors group, indicating that high expression of EEC cells, progenitor or differentiated, may be associated with a higher proportion of goblet cells, but were generally not pathologically defined as mucinous tumors (Supplemental Figure 1F & Supplemental Table 4).

The transcriptomic profile of maturing EECs has recently been elucidated using scRNA-seq, with robust expression markers identified at each stage of development (Gehart et al., 2019). Leveraging these markers, we inferred the potential stage of EEC progenitor maturation for each tumor sample. *NEUROG3*, one of the earliest markers of EEC differentiation, was significantly elevated in BRAF^EP^ compared to normal tissue and all other groups (Figure 1D & Supplemental Table 2). Generally, expression of early and intermediate markers was significantly higher in both BRAF^EP^ as well as KRAS^EP^ relative to all other tumor groups and significantly higher in BRAF^EP^ compared to KRAS^EP^. Conversely, expression of late markers *NEUROD1* and *LMX1A* were significantly lower in BRAF^EP^ than all other tumor groups. Consistent with lower expression of late EEC progenitor markers, markers of differentiated EECs were all significantly lower in BRAF^EP^ than in all other groups, with the exception of 4 out of 25 total comparisons where BRAF^EP^ expression was lower but did not reach significance (Figure 1E & Supplemental Table 2). These results suggest that differentiation of EECs may be halted at an early stage in some *BRAF* tumors.

### Organotypic cell lines recapitulate major populations of the large intestine

The HT29, NCI-H508 (H508), and LS174T organotypic cell lines have been used to extensively study the contribution of goblet cells to tumor progression. HT29 and H508 cells contain activating *BRAF* kinase domain mutations V600E and G596R, respectively, and LS174T cells contain the *KRAS* GTP-binding mutation G12D (Cancer Cell Line Encyclopedia)(Barretina et al., 2012). However, the presence and differentiation status of the EEC lineage in these cell lines is unknown. To further interrogate the secretory cell populations in these cell lines, we performed single-cell RNA sequencing (scRNA-seq) and compared our data against a select published scRNA-seq of the human colon (Supplemental Figure 2). Through marker analysis, it was determined that most major colon epithelial cell types were present in all three samples (Figure 2 A, B), including the secretory goblet and EEC populations (Figure 2C). The early secretory population was defined based on its varied transcriptional continuum, with genes such as *MEX3A* being expressed throughout the cluster, and other genes such as *DLL1* being uniquely expressed at the interface of the early EEC and goblet cell clusters (Supplemental Figure 3A, B). A combination of cell cycle-related gene expression profiling and marker analysis demonstrated that transit-amplifying cells (TAs) exist in the dataset in distinct cell cycle stages, consistent with previous literature and the cyclical-like shape of this population (Supplemental Figure 3C & 4). Paneth-like (PLC) and enterocyte-like cells (ELC) were also identified, as evidenced by markers such as *SPIB* and *LGALS2* respectively (Supplemental Figure 3D, E). However, well-defined markers for both populations were more clearly expressed in the normal colon, and marker expression was generally lower and more heterogeneous in the HT29 or H508 cell lines, indicating that these populations may not undergo canonical or complete differentiation in the cancer cell lines. We additionally identified a putative early goblet-like cell (EGLC) population that expressed some goblet cell markers such as *REG4*, *MUC5AC*, and *SERPIN1A* but was absent for *MUC2* (Figure 2A & Supplemental Figure 3F, *REG4*). Further, while intestinal stem-cell markers such as *LGR5*, *SMOC2*, and *OLFM4* were highly expressed in the normal colon, the expression of these markers was lower in H508 cells and virtually absent in HT29 cells. The absence of this population in our largest dataset (HT29) obfuscated the identification of a canonical stem population in our integrated analyses, however, these cells, which primarily originate from the colon sample, were positioned directly above our early secretory cluster (Supplemental Figure 3G, *SMOC2*).

**Figure 2:**
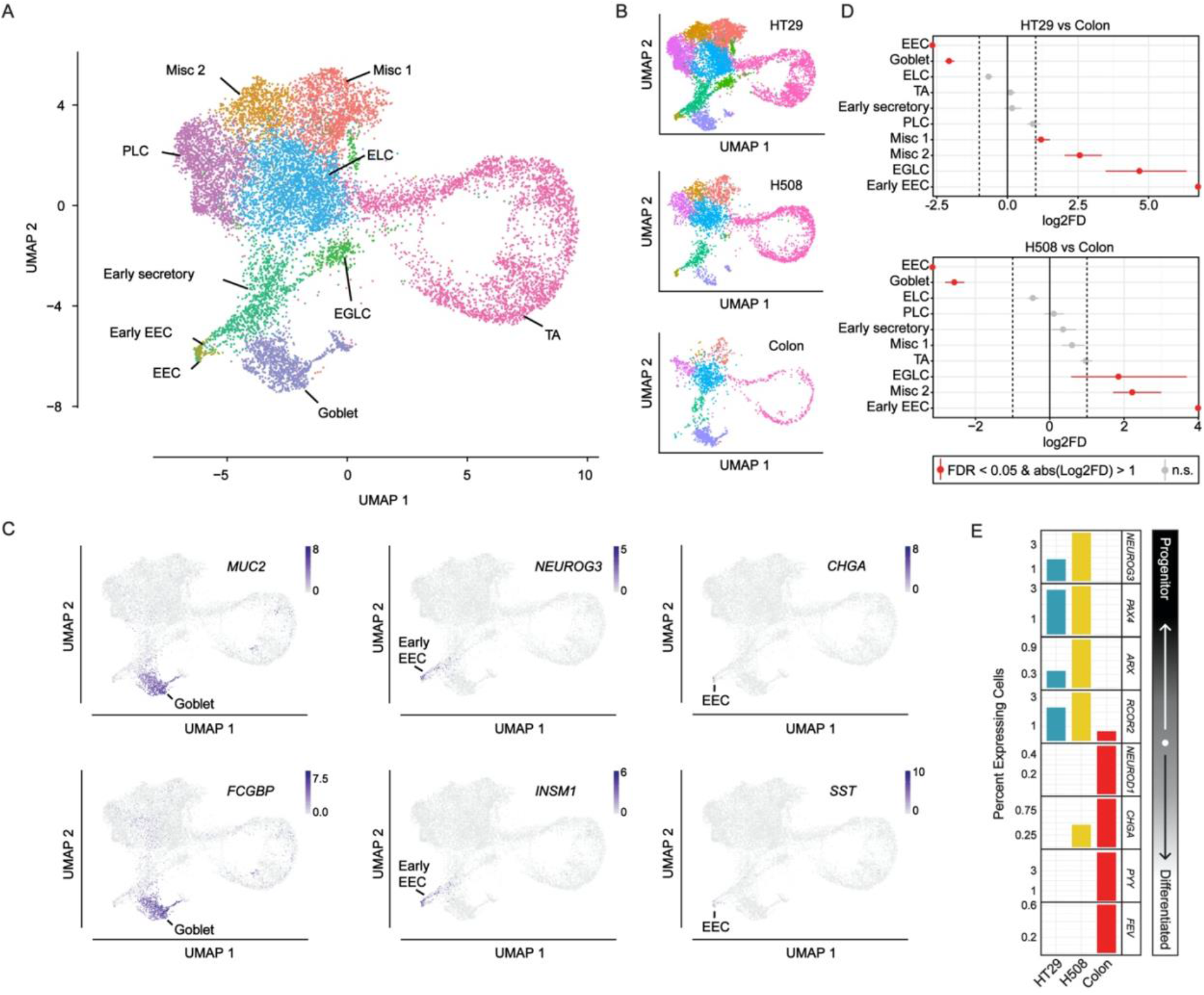
Multiple colon cell types are conserved in cell line models of CRC. A, Uniform Manifold Approximation and Projection for Dimension Reduction (UMAP) dot plot of a normal human colon, HT29 empty vector (EV), and H508 EV scRNA-seq samples colored by cell type/cluster. Cluster abbreviations: enteroendocrine (EEC), early goblet-like cells (EGLC), enterocyte-like cells (ELC), miscellaneous (misc 1 & misc 2), paneth-like cells (PLC), and transit-amplifying (TA). B, Individual UMAP dot plots of normal human colon, HT29 EV and H508 EV scRNA-seq samples colored by cell type/cluster. C, UMAP dot plots of normalized expression values of marker genes representative of the goblet, early EEC, and EEC cells in the combined colon, HT29 EV, and H508 EV samples. D, Relative differences in cell proportions for each cluster between the HT29 EV versus colon, and H508 EV versus colon. Clusters colored red have an FDR < 0.05 and mean | Log_2_ fold enrichment | > 1 compared to the normal colon (permutation test; n=10,000). E, Fraction of cells expressing various markers of EEC differentiation in the normal colon, HT29 EV, and H508 EV.

Compared to HT29 and H508 cells, the normal colon had significant enrichment of terminally differentiated secretory EEC and goblet cells (FDR < 0.05 and mean Log_2_ Fold Enrichment > 1) (Figure 2C). Conversely, the cell lines were enriched for early EECs as well as EGLCs and miscellaneous (Misc) cell clusters (FDR < 0.05 and mean Log_2_ Fold Enrichment > 1). The Misc clusters contained discrete markers, but these markers were not clearly associated with any one cell type (Supplemental Figure 3H, I). Since EGLCs were primarily present only in HT29 cells, we did not study this population further and rather focused on generally conserved cell types. The dichotomy between the cell lines and the normal intestine with respect to the differentiation status of EEC’s is echoed by examining the expression of well-established marker genes. While HT29 and H508 cells both express the early and intermediate EEC marker genes (*NEUROG3*, *PAX4*, *ARX*, and *RCOR2*), the normal colon only expresses intermediate and late markers (*RCOR2*, *NEUROD1*, *CHGA*, *PYY*, and *FEV*) (Figure 2D). Altogether these data indicate that some of the primary transcriptional regulators of secretory cell formation *in vivo* are present in the cell lines *in vitro*. Furthermore, and consistent with our findings from the TCGA, *BRAF* mutant cell lines are enriched for putative EEC progenitor cells compared to the normal colon.

### CpG methylation blocks *NEUROD1* expression in BRAF^EP^ cancers

We hypothesized that BRAF^EP^ tumors exhibit a molecular alteration that functions to halt EEC differentiation. The CpG island methylator phenotype (CIMP) is characterized by aberrant gains of methylation at dense regions of CpG nucleotides known as CpG islands (CpGI), and generally coincides with repressed gene transcription(Toyota et al., 1999). *KRAS* and *BRAF* mutant tumors are differentially associated with CIMP, with *BRAF-*mutant tumors being uniquely enriched for the CIMP high phenotype versus *KRAS* mutants(Weisenberger et al., 2006). Based on the significant difference in *NEUROD1* expression between BRAF^EP^ and all other tumors, we hypothesized that *NEUROD1* may be differentially methylated between these groups. Using TCGA COAD DNA methylation data, we determined both that methylation in all CRC groups was generally increased over the *NEUROD1* CpGI compared to normal, and that there was a strong negative correlation between CpGI DNA methylation and expression of *NEUROD1* (Figure 3A). In contrast, methylation was slightly reduced over the transcription start site of the non-CpGI containing gene PAX4 in all sample types and the correlation between methylation and PAX4 gene expression was weak or not present (Supplemental Figure 5). Among the tumor groups, BRAF^EP^ exhibited the highest levels of *NEUROD1* methylation while BRAF^EC^, KRAS^EP^, and KRAS^EC^ exhibited the lowest (Figure 3A), consistent with the lowest expression of differentiated EEC markers in BRAF^EP^ and highest expression in BRAF^EC^, KRAS^EP^, and KRAS^EC^ groups (Figure 1E). In line with the TCGA data, *NEUROD1* CpGI methylation was significantly higher in *BRAF* mutant HT29 and H508 cell lines compared to *KRAS* mutant LS174T cells or normal human colon organoids (Figure 3B). Treatment with a low dose of the DNA methyltransferase inhibitor decitabine (DAC) led to re-expression of *NEUROD1* in HT29 and H508 cell lines but did not change *NEUROD1* expression in LS174T cells, consistent with DNA methylation-dependent repression of *NEUROD1* in *BRAF*-mutant tumors (Figure 3C). To further show the specificity of DAC treatment-mediated reactivation of NEUROD1 in *BRAF*-mutant cells, we used ISX-9, which activates *NEUROD1* expression through a DNA-methylation independent mechanism(Schneider et al., 2008). Treatment with ISX-9 induced significant activation of *NEUROD1* in LS174T cells and *NEUROD1* re-expression in HT29 and H508 cell lines (Figure 3C). Co-treatment of DAC and ISX-9 further induced *NEUROD1* expression in HT29 and H508 cells, but not in LS174T cells. Further analysis revealed that *NEUROD1* promoter CpGI methylation was unchanged after ISX-9 treatment in HT29 cells, but significantly reduced after DAC treatment and reduced to a similar level with DAC/ISX-9 co-treatment (Figure 3D). Together this data suggests that BRAF^EP^ tumors exhibit CpGI methylation-dependent loss of *NEUROD1* expression, potentially contributing to the loss of EEC differentiation. Furthermore, HT29 and H508 cell lines may be appropriate models to study the regulation and function of EEC progenitors in colon cancer.

**Figure 3.**
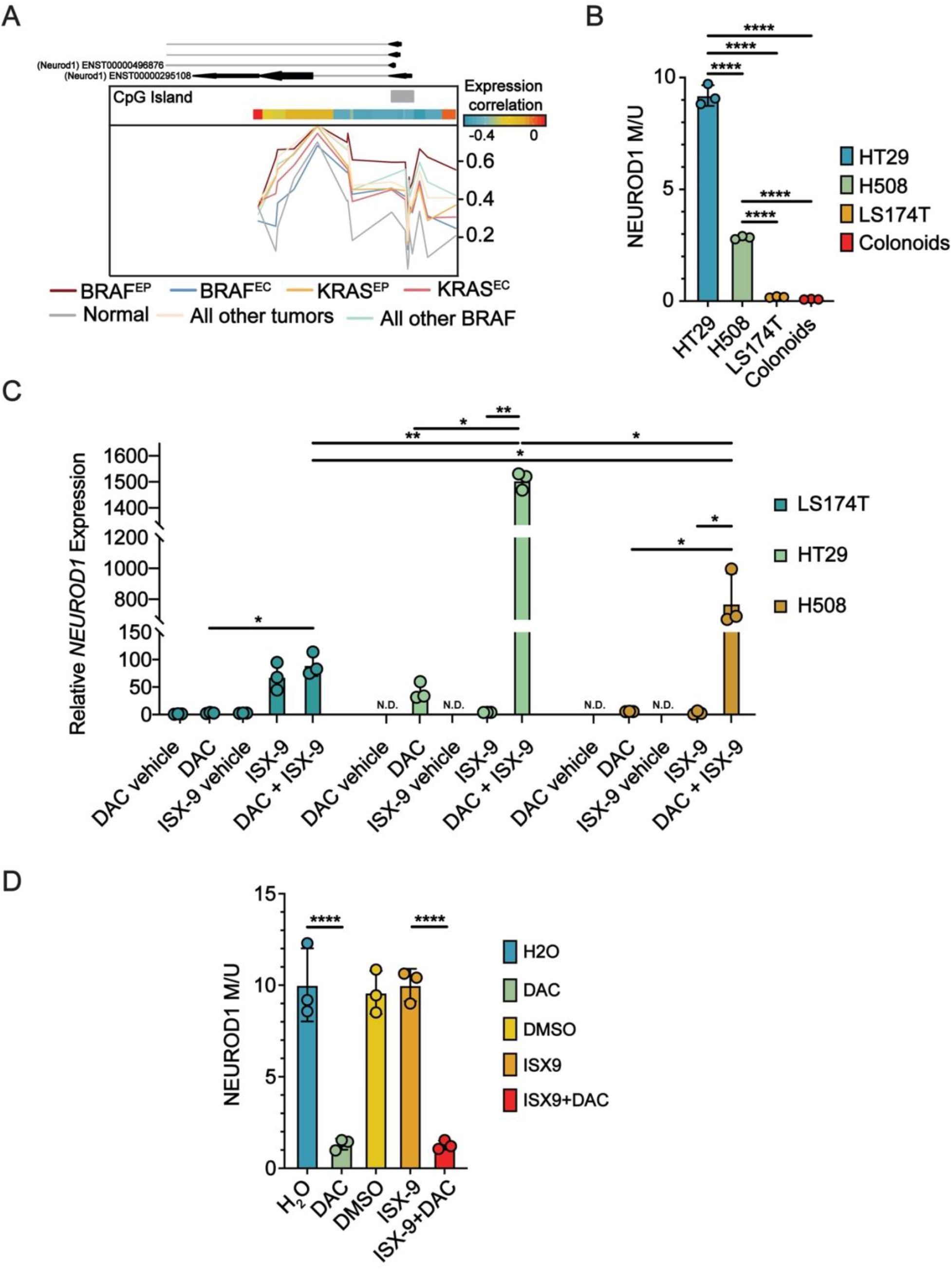
*NEUROD1* is silenced in BRAF^EP^ tumors via promoter CpG island methylation. A, Mean percent methylation from TCGA COAD Illumina human methylation 450K array probes across the transcription start site of *NEUROD1*. BRAF^EC^ (N = 5), BRAF^EP^ (N = 21), KRAS^EC^ (N = 23), KRAS^EP^ (N = 24), Normal (N = 41), All other BRAF (N = 33), All other Tumors (N = 329). The expression correlation heatmap depicts Spearman’s correlation between *NEUROD1* mRNA expression and 450K array methylation. B, Quantitative MSP at the *NEUROD1* promoter CpG island of bisulfite-treated DNA from normal colon organoids, LS174T, H508, and HT29 cells. C, LS174T, HT29, and H508 cells treated with H_2_O (DAC vehicle), 100 nM DAC for 72 hours, DMSO (ISX-9 vehicle), 40 μM ISX-9 for 24 hours, or a combination of DAC and ISX-9. *NEUROD1* expression is measured by RT-qPCR and normalized to housekeeping gene expression and LS174T DAC vehicle to establish baseline expression. D, Quantitative MSP at the *NEUROD1* promoter CpG island in bisulfite-treated DNA from HT29 cells treated as in B. Statistical analysis by (C) Welch’s t-test with Benjamini-Hochberg adjustment and by (B, D) one-way ANOVA with Tukey’s pairwise multiple comparisons testing (*, P <0.05; **, P <0.01; ****, P<0.0001).

### Trajectory analysis provides novel insight into the regulation of intestinal EEC and goblet cell commitment in CRC

The heterogeneous cell content of HT29 and H508 potentiates the use of computational trajectory analysis to better understand the transcriptomic profile of differentiating EECs and goblet cells in CRC. To this end, we performed RNA-velocity analysis, which uses the ratio of spliced to unspliced transcripts (analogous to mature versus nascent RNA) to infer cellular trajectories over developmental pseudotime. Furthermore, we extended the RNA-velocity analysis through integration with PAGA, which predicts the degree to which there is a connection between different clusters of cells based on the similarities and differences in their transcriptional profiles. For both the cell lines and colon samples an almost cyclical pattern emerges from within the TA cluster as seen in the velocity stream plots and the accompanying PAGA plots (Figure 4A). Within this cell type, velocity arrows can be seen flowing from the inferred S phase cells, through the G2/M phase cells, and finally into the G1 phase cells as would be expected (Supplemental Figure 3D). In the cell line samples, it can also be seen that the velocity stream bifurcates and flows either towards the goblet and EEC cell lineages, or towards the central cluster that contains PLCs, ELCs, and various other clusters (Figure 4A). Finally, CellRank was employed to further define clusters that are likely at an end state in differentiation. For both cell lines, the early EECs and goblet cells had a strong localized prediction to be terminally differentiated (Supplemental Figure 6). PLCs, and to some extent other surrounding cells in clusters such as ELCs, had a diffuse and less well-localized prediction as terminally differentiated. Additionally, a subset of cells in TA clusters that varied between samples was predicted to represent an end state, likely reflecting cells that continue to cycle. It is noted that in the colon sample, the disconnected cell populations and low relative cell and sequencing read depth resulted in less informative RNA-velocity and CellRank results. Whereas for the cell lines the connectedness of clusters and relatively high depth of coverage leads to more continuous and confident velocity streams, and therefore more informative downstream analysis such as CellRank.

**Figure 4:**
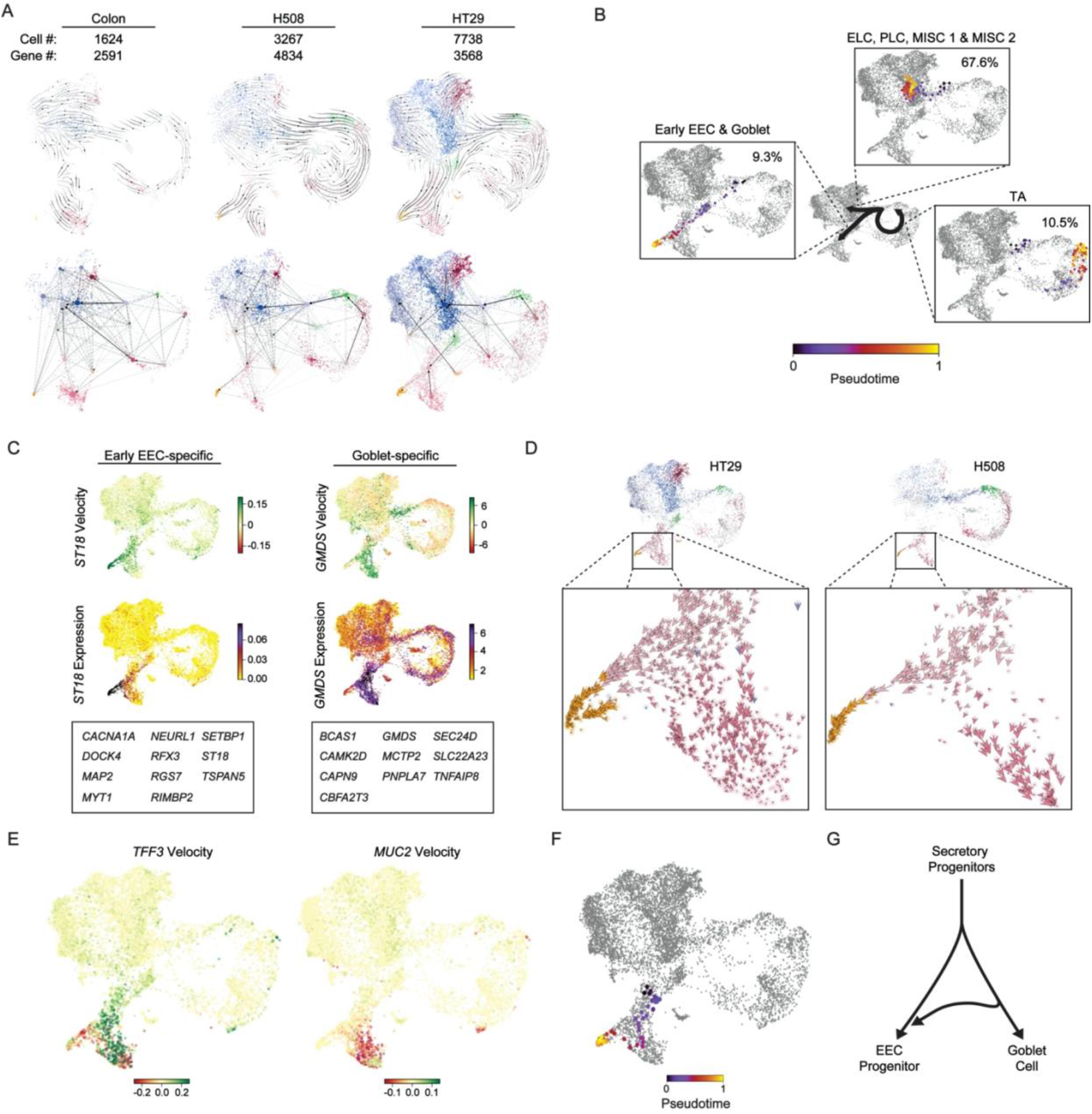
Integrating high-resolution single-cell datasets provides detailed insights into the differentiation of distinct intestinal cell lineages. A, RNA velocity stream plots (top) and directed PAGA plots (bottom) of the colon, H508 EV, and HT29 EV scRNA-seq data colored by clusters from spliced reads, showing inferred differentiation trajectories. B, Representative plots of inferred cell descendants over pseudotime from TA cells in G2/M phase, and the approximate percentages of their inferred end-state cell type in HT29. C, Velocity and expression UMAP plots, and a table of select markers that have differential velocity in the goblet or early EEC cells versus all other clusters in HT29 and H508. D, Velocity arrows for individual cells in the early secretory, goblet, and early EEC cell clusters. E, UMAP plot of RNA velocities of *TFF3* and *MUC2* per cell in HT29 and H508. F, Representative descendent tracing through pseudotime of a cell originating in the early secretory cluster, and that is inferred to slightly differentiate towards the goblet cell lineage before becoming an early EEC in HT29. G, Diagram depicting velocity analysis-based model of secretory cell differentiation from early secretory progenitors toward early EEC and goblet cells, as well as the redirection of some cells away from terminal goblet cell differentiation toward early EECs.

Using the RNA-velocity results and the knowledge of inferred cell end states from CellRank, we explored the goblet and EEC cell lineages in our cell lines by predicting the “descendants” of cells, or the cells they are most likely to become moving forward in pseudotime/differentiation. Using this feature, we inferred the descendants of TA cells at the end of the G2/M phase as they potentially exit the cell cycle and begin specification toward CellRank predicted end-states. Approximately 10% of these cells had descendants that were predicted to be in the goblet and EEC lineages (Figure 4B), consistent with the relatively small proportion of these cells compared to the whole cellular population. Approximately 80% of cells were predicted to transition into other cell types such as ELC and PLCs or continued cycling, and the remaining 10% did not have well-defined descendants. Using the ratio of spliced to unspliced reads for a gene in a cell, RNA-velocity can also infer the gene velocity, or whether a gene is likely being activated or repressed in a subpopulation of cells. If the proportion of unspliced reads increases, activation (high velocity) is assumed since more nascent transcripts would be expected to be captured by a gene that is transcribing at a higher rate, and vice versa for repression. Leveraging this information, we looked for genes with uniquely high or low velocities in our goblet and EEC clusters to better understand the regulatory landscape of their transcription. High confidence targets identified through this analysis include genes such as *MYT1*, which has previously been shown to function as a general specification factor for the EEC lineage(Gehart et al., 2019) (Figure 4C). Other high-velocity genes, *RFX3* and *ST18*, have been implicated in the formation and function of pancreatic endocrine cells, and may therefore also function similarly in the large intestine(Ait-Lounis et al., 2010) (Henry et al., 2014). In addition to early EEC lineage specifiers, we identified genes associated with cell morphology like *MAP2*, the product of which is involved in the formation of dendrites(Harada et al., 2002). As evidence of this function *in vitro*, analysis of early EEC morphology by immunofluorescence shows that these cells exhibit large ramified processes reminiscent of the dendrite pseudopod structures of mature EECs, with a terminal synaptic-like bouton(Supplemental Figure 7A). With respect to goblet cells, *CBFA2T3* has an established role in the specification of these cells(Amann et al., 2005) (Figure 4C). High-velocity genes toward the goblet population were generally involved in calcium signaling and metabolism, such as *GMDS*. In contrast to the early EECs, goblet cells primarily exhibited ovoid-like morphologies with some cells exhibiting the classic signet-ring morphology with nuclei pressed against the cell membrane (Supplemental Figure 7A).

Individual cell velocity arrows indicated that cells at the furthest points of the early EEC and goblet cell clusters had greatly reduced velocity, which means they are predicted to be terminally differentiated (Figure 4D & Supplemental Figure 7B). For the goblet cell cluster, the pattern of predicted activation or repression of certain genes appears to correspond to this boundary. For example, *Tre-foil factor 3* (*TFF3)* appears to switch from a predicted activated state to a repressed state near the terminally differentiated cells, while *MUC2* goes from a repressed to activated state (Figure 4E). As previously described, the secretory cell lineage has an interesting bifurcating shape, potentially due to a fate decision for common progenitors. Zooming into that region and looking at the velocity arrow direction and speed for individual cells reveals a strong bias for many cells towards the EEC lineage in the cancer cell lines (Figure 4D). In juxtaposition, the directionality and speed of cells differentiating to goblet cells is less coherent, with many cells seeming to change direction back towards the undifferentiated or EEC lineages. A representative descendent plot of an early secretory cell predicted to differentiate towards the goblet cell fate, and then redirect towards the early EEC fate is shown (Figure 4F).

To directly test the apparent shift from goblet cell differentiation toward early EEC seen in these cell lines, genetic knockdowns of early EEC specification gene *NEUROG3* and goblet cell specification gene *SPDEF* were generated. *SPDEF* KD reduced *NEUROG3* and *MUC2* expression, while *NEUROG3* KD had no effect on *SPDEF* or *MUC2* expression (Supplemental Figure 7C). These observations suggest a potential model in these CRC cell lines where some cells that are otherwise poised to form goblet cells are redirected prior to terminal differentiation to become early EECs (Figure 4G).

### Secretory cells and their progenitors secrete tumor-associated proteins in CRC

Secretory progenitors exist in highly transient states so are rarely captured in the normal intestine, and are therefore difficult to study. Consistent with our TCGA expression data, the proportion of mature *ATOH1* expressing cells was higher in the colon than in HT29 or H508 cells, where the numbers of early EEC *NEUROG3* expressing cells are higher in the CRC cell lines (Supplemental Figure 8). While present in these cell lines, *NEUROG3* expressing cells nonetheless make up only ∼2% of the total cell line population. The enrichment of these early EEC progenitors in the cell lines provides a unique opportunity to study in cancer the functions and interactions between these otherwise highly transient populations.

To interrogate the secretory protein landscape mediated by early EECs, EECs, and goblet cells, the top 50 markers for each of these cell types were cross-referenced to the Human Protein Atlas (proteinatlas.org) as well as related literature, to identify secreted proteins. All populations expressed various secreted factor genes, many of which have been shown to enhance cancer-associated phenotypes (Figure 5A; Denoted in red). For example, early EECs express *ENPP2*, which has been shown to promote cell migration. *TFF3* is canonically considered a goblet cell-secreted factor that displaces an antagonist of the EGFR, thereby enabling enhanced activation of the PI3K-AKT pathway(Belle et al., 2019). Uniquely, *TFF3* is highly expressed in all three secretory cell types. Our trajectory analysis indicates that *TFF3* gains velocity in stable EEC progenitors, where it is highly expressed (Figures 4E & 5A). Further analysis indicates that *TFF3* is generally highly expressed in goblet and EEC lineage cells compared to all other groups (Figure 5B). Genetic knockdown of *ATOH1* or *NEUROG3* to decrease secretory and EEC populations, respectively, lead to significant loss of *TFF3* expression (Figure 5C, D & Supplemental Figure 7C) and significantly less TFF3 secretion into the media (Figure 5E). Consistent with the previously described role for TFF3 in promoting cell survival(Taupin et al., 2000), the addition of a neutralizing TFF3 antibody to serum-free media resulted in a significant reduction in cell survival during EGF signaling (Figure 5F). These data provide evidence that EEC progenitors produce secreted peptides that may contribute to tumor progression, such as TFF3, which is critical for cell survival during growth factor signaling.

**Figure 5:**
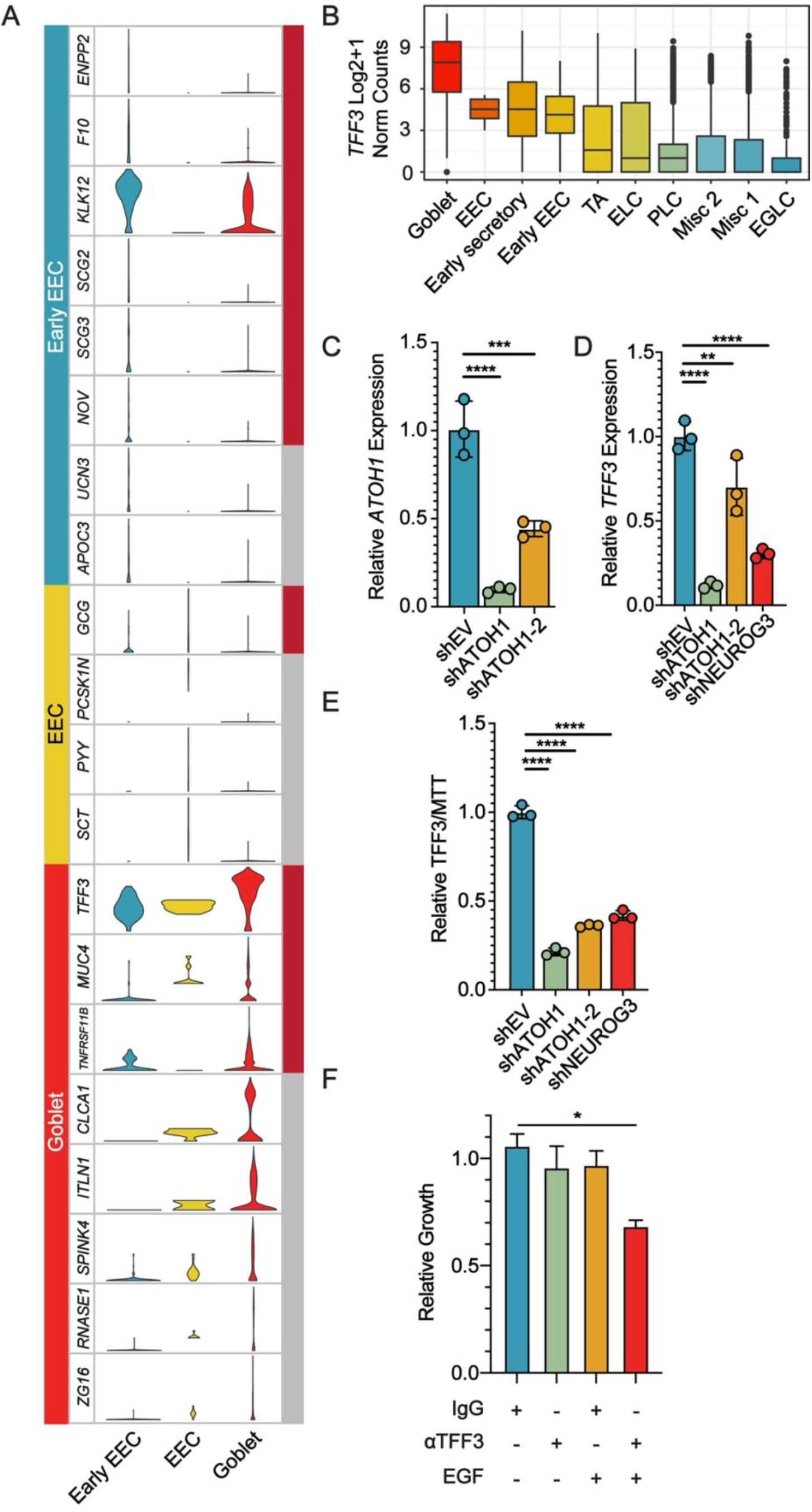
Secretory cells and their progenitors secrete tumor-promoting factors. A, Violin plots of normalized Log_2_ + 1 expression for various marker genes in early EEC, EEC, and goblet subpopulations from the combined scRNA-seq data. Each gene is scaled separately. The far-right column denotes the tumor-promoting status of each gene, with dark red boxes indicating an association with tumor progression, and grey indicating either tumor-suppressive or no published link to tumorigenesis. B, Normalized expression of *TFF3* in cell subpopulations of the colon, HT29, and H508 scRNA-seq datasets. Boxplot includes the median and box edges represent 2nd and 3rd quartiles and whiskers represent 1st and 4th quartiles. C, RT-qPCR of *ATOH1* RNA expression level in EV (shEV) and ATOH1 KD (shATOH1) cells. D, RT-qPCR of *TFF3* RNA levels in HT29 cells after the indicated knockdown. E, ELISA of TFF3 protein levels in media, normalized to MTT growth assay and EV. F, 24-hour cell growth determined by the CellTiter-Glo Luminescent Cell Viability Assay. Cells were treated with 1 μg/ml αTFF3 or IgG, alone or in combination with 125 ng/ml EGF. C, D, & E Statistical analysis by one-way ANOVA with Dunnett’s or (F) Tukey’s pairwise multiple comparisons testing (*, P <0.05; **, P <0.01; ***, P<0.001; ****, P<0.0001).

### LSD1 knockdown results in loss of secretory cells

To better understand the regulation of EECs in CRC, we performed pathway enrichment analysis of differentially expressed genes between the early EECs and goblet cells, using the Reactome Pathway Database(Jassal et al., 2020). Compared to goblet cells, early EECs were significantly enriched for chromatin related terms such as “Chromatin modifying enzymes”, and “Chromatin organization” (FDR < 0.05) (Supplemental Figure 9). The most differentially expressed chromatin-modifying enzyme was *Lysine-specific demethylase 1* (*LSD1*) (FDR of 2.97×10^−14^ and Log_2_ FC of 1.03). Generally, *LSD1* expression was highest in early EEC and early secretory cells, lowest in differentiated EEC and goblet cells, and similar in all other clusters (Supplemental Figure 10A). We have previously demonstrated that loss of LSD1 significantly alters clonogenic growth in organotypic HT29 cells, but not in stem-like SW480 cells, which do not form secretory cells(Miller et al., 2020). Similarly, reduction of EEC or secretory cells by knockdown of *NEUROG3* or *ATOH1* significantly reduced clonogenic growth in HT29 cells (Supplemental Figure 10B). Combining this observation with our *LSD1* expression analysis, we hypothesized that *LSD1* expression levels increase in secretory progenitors to regulate EEC progenitor and goblet cell formation in our cell lines. Additionally, loss of these populations after LSD1 KD may be responsible for our previously reported clonogenic growth phenotypes(Miller et al., 2020).

As a preliminary analysis to determine if LSD1 may be necessary for the formation of secretory lineage cells, we reanalyzed our previously published LSD1 knockdown (KD) RNA-seq data in HT29 and SW480 cells(Miller et al., 2020). LSD1 KD was associated with a significant reduction of nearly all tested EEC progenitor and goblet cell markers in HT29 cells, but no markers were reduced in SW480 cells (FDR < 0.05) (Supplemental Figure 10C). To deconvolute whether we lose these cell populations or have a global decrease in transcript expression for these genes, we performed scRNA-seq on HT29 and H508 cells after LSD1 KD and integrated these samples into the previously analyzed empty vector (EV) controls using Seurat, as described in the Supplemental Methods. Consistent with reports of *LSD1* overexpression in CRC(Hsu et al., 2015), *LSD1* expression was elevated in our cell lines compared to the normal colon, which had expression levels similar to the HT29 and H508 LSD1 KD samples (Supplemental Figure 10D). Loss of LSD1 dramatically reduced relative cell-type proportions of early EEC and goblet cells (mean Log_2_ Fold Difference < −1 and FDR < 0.05) but did not significantly decrease other populations in both cell lines (Figure 6A). To better understand the effect of LSD1 KD on the differentiation trajectory of subpopulations within the cell lines, we computed the RNA-velocity and compared the results to the EV controls. Strong relative velocity length and confidence can be observed in the goblet and EEC lineages in the controls (Figure 6B). Velocity confidence is a measure of agreement in the directionality of differentiation between nearby cells, and velocity length is a measure of the relative “speed” that one subpopulation is differentiating towards another in pseudotime. In LSD1 KD cells, however, both velocity length and confidence toward the secretory lineage is diminished relative to the other non-secretory cell lineage clusters such as TAs. As validation of these results, LSD1 KD leads to a significant reduction in the percentage of MUC2 positive goblet cells and SUSD2 positive cells, a marker expressed in EEC progenitor populations, as determined by immunofluorescence (Figure 6C, D). Further, while LSD1 KD reduced expression of *NEUROG3* and *ATOH1* in HT29 and H508 cells, this reduction was not observed in *KRAS* mutant LS174T cells (Supplemental Figure 10E).

**Figure 6:**
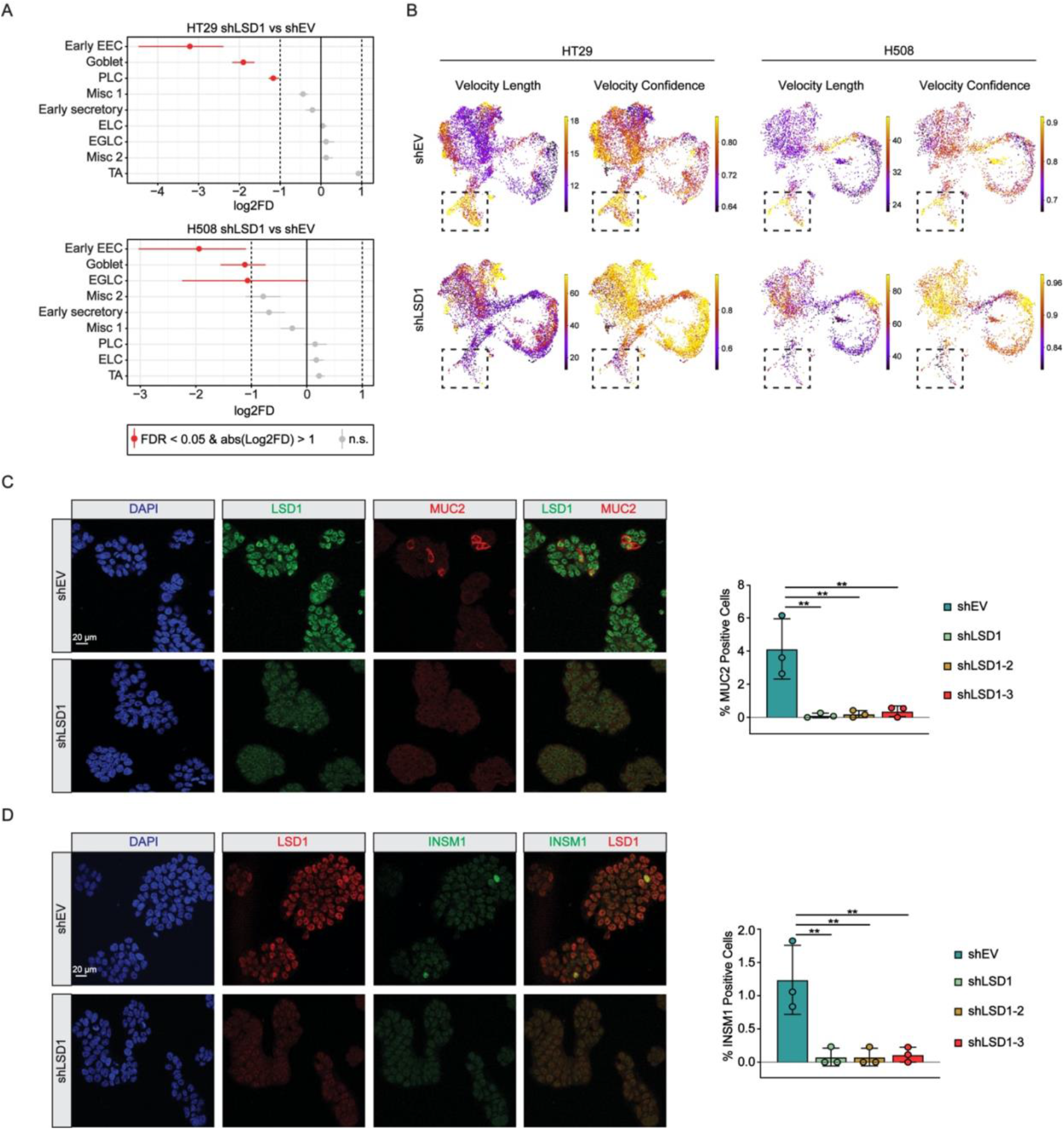
LSD1 is required for the persistence of enteroendocrine progenitor and goblet cells in *BRAF* mutant CRC. A, Relative differences in cell proportions for each cluster between the HT29 LSD1 KD (shLSD1) versus EV (shEV), and H508 LSD1 KD versus EV. Clusters colored red have an FDR < 0.05 and mean | Log_2_ fold enrichment | > 1 compared to normal colon (permutation test; n=10,000). B, UMAP plot showing RNA velocity length and confidence in HT29 EV and LSD1 KD as well as H508 EV and LSD1 KD. The dotted box indicates the region of interest, including part of the early secretory cluster and the entirety of the early EEC and goblet clusters. Representative immunofluorescence images of EV or LSD1 KD cells depicting (C) MUC2 signal and quantification of percent positive cells or (D) INSM1 signal and quantification of percent positive cells. Statistical analysis by one-way ANOVA with Dunnett’s pairwise multiple comparisons testing (**, P <0.01).

Disruption of the NOTCH pathway can significantly change the relative levels of secretory cells, and in HT29 cells NOTCH inhibition by DBZ significantly increased secretory cells at the expense of HES1+ absorptive cells as inferred by changes in gene expression (Figure 1A & Supplemental Figure 11A). LSD1 KD did not alter *HES1* expression in any cell line, and further analysis of our previously published RNA-seq data shows that LSD1 KD did not alter the notch signaling pathway, indicating that LSD1 function may be independent of NOTCH1 specification (Supplemental Figure 10E, 11B). Together these data show that LSD1 is required for the formation of secretory EEC progenitors and goblet cells in HT29 and H508 cell lines.

### Loss of secretory cells reduces pS473-AKT and suppresses tumor growth and metastasis

We have previously demonstrated that LSD1 regulates phosphorylation of AKT at serine 473, a modification involved in the activation of AKT through an unknown mechanism(Miller et al., 2020). We hypothesized that the loss of secretory cells following LSD1 KD may thereby abrogate AKT phosphorylation potentially due to loss of secreted factors. Replacing the media of LSD1 KD cells with media conditioned by other LSD1 KD cells resulted in a significant decrease in pS473-AKT compared to EV cells (Figure 7A). Replacing the media of LSD1 KD cells with media conditioned by EV cells resulted in the significant rescue of pS473-AKT levels and increased phosphorylation of the AKT-downstream target TSC complex subunit 2 (TSC2). These results suggest that LSD1 KD cells are unable to produce a secreted factor critical for the activation of AKT, likely due to the loss of the cell type secreting this factor.

**Figure 7:**
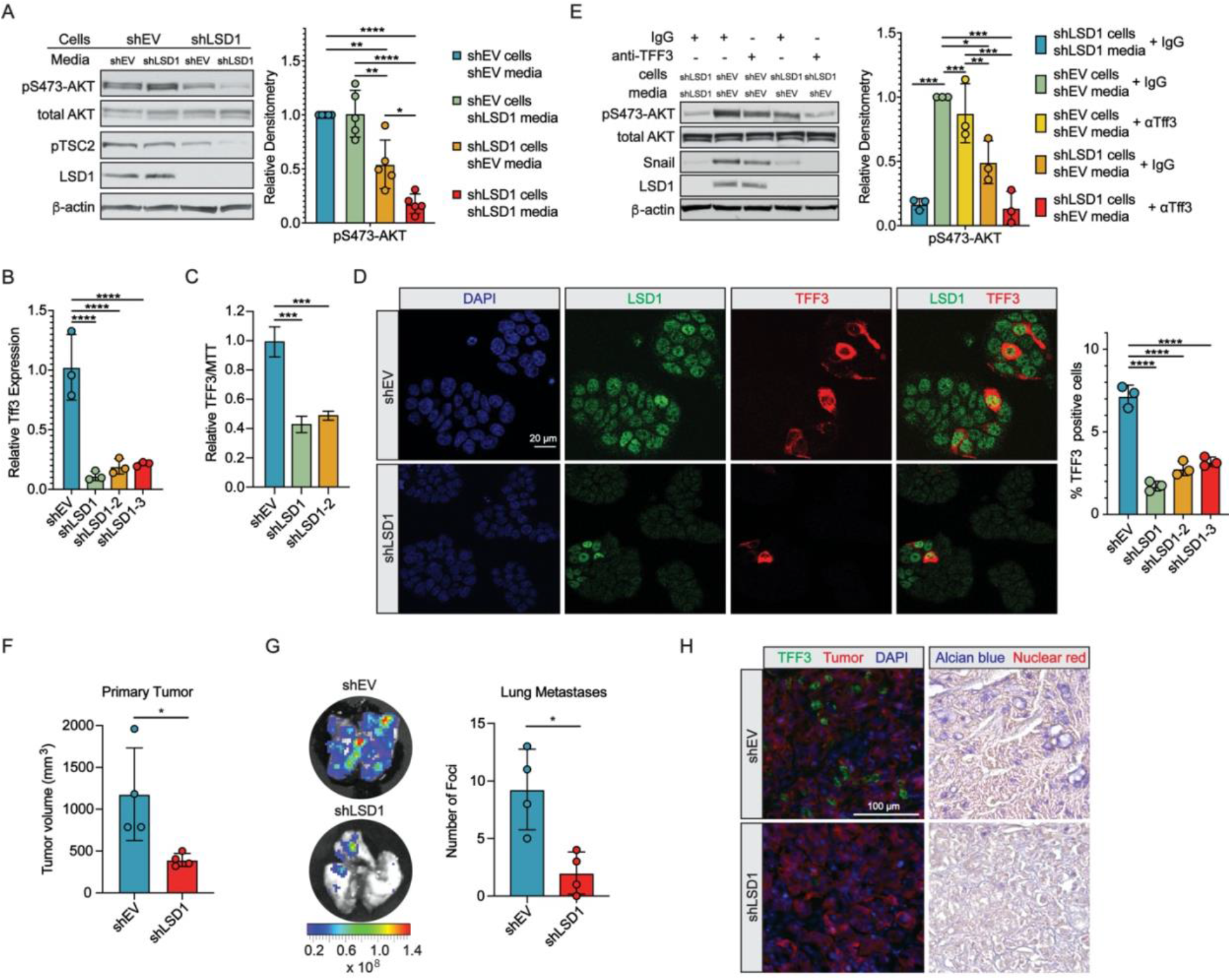
LSD1 facilitates AKT activation and tumor progression by promoting the formation of secretory cells. A, Western blot of EV (shEV) or LSD1 KD (shLSD1) HT29 cells following media transfer (n=5). Western blots quantified by densitometric analysis and normalized to β-actin and EV control. B, RT-qPCR of *TFF3* RNA expression levels after LSD1 KD with 3 different shRNA. C, ELISA of TFF3 protein levels in media, normalized to MTT growth assay and EV. D, Representative immunofluorescence images of EV or LSD1 KD cells and quantification of percent positive cells from three independent shRNA targeting LSD1. E, Western blot of EV or LSD1 KD cells following media transfer in combination with IgG or αTFF3. Western blots quantified by densitometric analysis and normalized to β-actin and EV control. F, Measurement of primary tumor volume or (G) number of lung metastatic foci from HT29 orthotopic xenograft NOD SCID mice (n=4). Scale represents total flux (photons/sec/cm2). H, Immunofluorescence, and brightfield images of frozen primary xenograft tumors. Images are representative of two biological replicates. A & E, Statistical analysis by one-way ANOVA with Tukey’s pairwise multiple comparisons testing and (B-D) Dunnett’s multiple comparison testing, and (G,H) by Welch’s t-test (*, P <0.05; **, P <0.01; ***,P<0.001; ****, P<0.0001).

LSD1 KD significantly reduced *TFF3* mRNA expression, TFF3 protein secretion into the media, and TFF3 positive cells (Figure 7B-D). Given the crucial function of TFF3 protein in the activation of EGFR, we tested the hypothesis that loss of TFF3 production in LSD1 KD cells reduces pS473-AKT. Treatment with a TFF3-neutralizing antibody reduced basal pS473-AKT and completely blocked pS473-AKT rescue in LSD1 KD cells supplemented with EV media (Figure 7E). SNAIL protein levels followed a similar trend, in agreement with our previous work demonstrating that LSD1 regulates SNAIL protein stability through the regulation of AKT activation(Miller et al., 2020).

To elucidate the contribution of LSD1-mediated specification of secretory cells to CRC *in vivo*, HT29 cells dual-labeled with luciferase and mCherry were orthotopically implanted into the mouse colon. LSD1 KD tumors exhibited significantly reduced volume at the primary site and significantly reduced metastases to the lung (Figure 7F, G). LSD1 KD tumors exhibited reduced staining for TFF3 and mucinous vacuoles (Figure 7H). Furthermore, while the mature EEC marker Synaptophysin was detected in the mouse large intestine, no positive staining was detected in mCherry tumor cells, consistent with the lack of EEC differentiation observed *in vitro* (Supplemental Figure 12). Together these data demonstrate that loss of secretory cells following LSD1 KD leads to loss of pS473-AKT and abrogates tumor growth and metastases *in vivo*.

## Discussion

*BRAF* mutant CRCs can be highly aggressive, with many studies showing worse survival compared to other CRC subtypes(Yaeger et al., 2014) (Margonis et al., 2018). In general, therapy response has been variable in clinical trials targeting patients with *BRAF* mutations, highlighting the need to develop additional therapeutic targets(De Roock et al., 2010) (Yaeger et al., 2015). In this study, we demonstrate that in addition to goblet cells, a significant proportion of *BRAF* mutant CRCs are also enriched for EEC progenitors, and support the notion that the presence of secretory cell populations is an important consideration in the treatment of *BRAF* mutant CRC. Furthermore, we identify LSD1 as a potential druggable target in secretory-cell-enriched *BRAF* mutant CRC. BRAF mutant CRC is strongly associated with CIMP(Weisenberger et al., 2006). Currently investigated treatments for *BRAF* CRC include DNA methyltransferase inhibitors (DNMTi) such as 5-azacitidine, which has been shown to sensitize *BRAF* mutant CRC cell lines to BRAF inhibitors(Mao et al., 2013). Interestingly we find that hypermethylation of the *NEUROD1* promoter CpGI is associated with a differentiation block in EECs resulting in an accumulation of EEC progenitors in *BRAF* mutant CRC. Treatment with the DNMTi decitabine demethylates the promoter CpGI resulting in re-expression of low levels of *NEUROD1* and the ability of neurogenesis-inducing small molecule ISX-9 to induce robust expression of *NEUROD1* when used in combination with decitabine. These findings suggest that EEC progenitors in *BRAF* mutant CRC can be further differentiated into EECs through the use of DNMTis. Beyond our findings in CRC, *NEUROD1* is frequently differentially methylated in neoplastic breast tissue(Fiegl et al., 2008). While the function of this methylation is unclear, these patients have a much worse overall survival, indicating a potential molecular connection between these two cancer types. Furthermore, *NEUROD1* is repressed by the polycomb repressive complex 2 (PRC2) in medulloblastomas as a mechanism to stymie tumor cell differentiation(Cheng et al., 2020). The PRC2 has been shown to mark genes for *de novo* DNA methylation in multiple cancer types including CRC suggesting that the PRC2 may be linked to *NEUROD1* CpGI methylation in BRAF-mutant CRC(Schlesinger et al., 2007).

The transient nature of EEC progenitors makes these populations more difficult to study than their fully differentiated counterparts, which still represent less than 1% of the adult large intestine. Our finding that *BRAF* mutant CRC cell lines maintain high levels of EEC progenitors suggests the cell lines may provide suitable models for the study of secretory cells in progenitor states. When comparing our scRNA-seq data from HT29 and H508 cell lines to a normal human colon sample, multiple of the clearly defined cell types seen in the colon sample appeared in a more undifferentiated or transcriptionally altered state in the CRC cell lines, consistent with other studies of CRC tumors at single-cell resolution(Uhlitz et al., 2020). However, distinct cell types such as goblet cells and putative early EECs appeared well-defined and with clearly conserved markers in both the normal colon and CRC cell lines. In line with these findings, researchers have derived subclones from the HT29 cell line capable of forming polarized goblet and enterocyte monolayers(Huet et al., 1987). RNA velocity predicted a pattern of cell differentiation for these populations where a subset of TA cells presumably finishing the cell cycle differentiate towards early progenitors and bifurcate into the stable goblet and early EEC cells. In our analysis of this data, we identify recently established transcriptional drivers of EEC and goblet cell differentiation as well as potential novel regulators. Due to the clear conservation of core transcriptional profiles in secretory populations between our *BRAF* mutant CRC cell lines and the normal colon, it is likely that most of our findings are applicable and of interest to the study of normal differentiation in the colon.

One of the clusters that were variant in the CRC cell lines compared to the normal intestine was canonical stem cells, which were completely absent in HT29 cells. We note in our data that in HT29 cells *CDX2* expression was nearly absent, the loss of which frequently occurs in *BRAF* mutant CRC, and contributes to sessile serrated adenoma progression, a tumor subtype that progresses rapidly(Sakamoto et al., 2017). *CDX2* is required to maintain the identity of intestinal stem cells(Simmini et al., 2014) and therefore the lack of *CDX2* expression in HT29 may be linked to the lack of canonical stem cell populations, and explain the absence of this population in this cell line. Another interesting observation more specific to the CRC cell lines was the presence of the EGLC cluster. This cluster contained some canonical markers of goblet cells but was predicted based on the trajectory analyses to be a transition state in the formation of differentiated secretory cells. Given the deeper sequencing of the HT29 cell line, it is unclear whether or not this population was identified due to the resolution of this dataset or whether it may be due to the underlying biology of this specific cancer.

Further analysis of EEC progenitors in our cell lines showed that these cells surprisingly secrete factors such as TFF3, a factor previously thought to be secreted predominantly just by goblet cells. We demonstrate that TFF3 promotes cell survival during growth factor signaling suggesting both EECs and goblet cells secrete factors that promote CRC. Based on our identification of other tumor-promoting secretory genes expressed by goblet cells and EECs, other secreted factors in addition to TFF3 also likely contribute to CRC progression. Furthermore, our data also suggest EEC progenitors are important in colony formation, a critical step in metastasis. In support of this notion, another study demonstrated that NEUROG3+ cells not only give rise to EECs but surprisingly also to non-EEC cells, which suggests these progenitors can dedifferentiate to produce other lineages(Schonhoff et al., 2004). Additional recent studies have shown that ATOH1+ common secretory progenitors can dedifferentiate into a multipotent stem-like state spontaneously and in response to intestinal injury(Tomic et al., 2018) (Castillo-Azofeifa et al., 2019). In agreement with these studies our trajectory analyses predict that while the vast majority of EEC progenitors are stable, a small fraction is predicted to dedifferentiate. It is tempting to speculate that the dedifferentiation of secretory progenitors may play a role in successful colony formation during metastases of *BRAF* tumors.

We demonstrate for the first time that LSD1 is required for the maintenance of secretory EECs and goblet cells in *BRAF* CRC. By employing scRNA-sequencing we were able to determine that LS1 KD leads to the loss of velocity of lineage specifying genes in early secretory cells, culminating in the loss of early EEC and goblet terminal secretory states. Interestingly, we determined that LSD1 KD did not alter secretory cells in the *KRAS* mutant mucinous LS174T CRC cells. This study sheds light on previous work demonstrating that LSD1 regulates activation of AKT in HT29 cells (Miller et al., 2020) as we now show secreted TFF3 may play a critical role in this process. Reduction of goblet and EEC cells by LSD1 KD resulted in reduced TFF3 secretion, allowing LSD1 KD to ablate AKT activation in the whole cell population even though it is only directly affecting a small percentage of cells. We find that LSD1 appears to act downstream of NOTCH signaling during the differentiation process. We further show that LSD1 levels are highest in EEC progenitors and early secretory cells. In accordance, we speculate that LSD1 may function directly in early secretory progenitors, to regulate the expression of genes important for the formation of further differentiated EEC progenitors and goblet cells. In vivo, LSD1 KD xenografts maintained the loss of secretory cells seen in cell culture and reduced tumor growth and metastasis. Future studies that establish a mechanistic understanding of how LSD1 regulates the maintenance of secretory cells will be critical to translate these findings into rational therapeutic options for patients with *BRAF* mutant CRC.

## Author Contributions

Conceptualization, S.A.M., R.A.P., and H.M.O.; Methodologies, S.A.M., R.A.P., S.S., T.L.,T.D.H., C.A.L, and H.M.O.; Computational Analysis, S.A.M., R.A.P., and T.L.; Writing, S.A.M., R.A.P., H.M.O., and G.E.Z.; Supervision, H.M.O. and G.E.Z.; Funding Acquisition, H.M.O. and S.A.M.

## Acknowledgments and Funding

We thank the Indiana University Light Microscopy Imaging Center and the Center for Medical Genomics at Indiana University School of Medicine, which is partially supported by the Indiana University Grand Challenges Precision Health Initiative. We would additionally like to thank Dr. Roberto Pili at the Indiana University School of Medicine for generously providing laboratory space to Samuel A Miller to prepare samples for scRNA-sequencing. We also thank Sue Childress for her assistance with tissue processing. This work was funded, in part, with a Core Pilot Grant [to H.M. O’Hagan] from the Indiana Clinical and Translational Sciences Institute funded, in part by Grant Number ULITR002529 from the National Institutes of Health, National Center for Advancing Translational Science, Clinical and Translational Sciences Award. Support was also provided by the National Institutes of Health, National Center for Advancing Translational Sciences, Clinical and Translational Sciences Award [TL1 TR001107 and UL1 TR001108 (principal investigator, A. Shekar), to S.A. Miller]. This work was also supported by a Research Enhancement Grant and Elwert Award [to H. M. O’Hagan] from the Indiana University School of Medicine. The content is solely the responsibility of the authors and does not necessarily represent the official views of the National Institutes of Health or the Indiana University School of Medicine. Additional research funding was provided by Van Andel Institute through the Van Andel Institute - Stand Up to Cancer Epigenetics Dream Team. Stand Up to Cancer is a division of the Entertainment Industry Foundation, administered by AACR.

## References

Abdalla, A.S., Khan, K.A., Shah, A., Asaad, A., Salter, V., Barron, M., Eldruki, S., Salih, V., and Alowami, S.O. (2020). Colonic Goblet Cell Carcinoid: Rarity of a Rarity! A Case Report and Review of Literature. Chirurgia (Bucur) 115, 102–111.

Ait-Lounis, A., Bonal, C., Seguin-Estevez, Q., Schmid, C.D., Bucher, P., Herrera, P.L., Durand, B., Meda, P., and Reith, W. (2010). The transcription factor Rfx3 regulates beta-cell differentiation, function, and glucokinase expression. Diabetes 59, 1674–1685.

Amann, J.M., Chyla, B.J., Ellis, T.C., Martinez, A., Moore, A.C., Franklin, J.L., McGhee, L., Meyers, S., Ohm, J.E., Luce, K.S., et al. (2005). Mtgr1 is a transcriptional corepressor that is required for maintenance of the secretory cell lineage in the small intestine. Mol Cell Biol 25, 9576–9585.

Barretina, J., Caponigro, G., Stransky, N., Venkatesan, K., Margolin, A.A., Kim, S., Wilson, C.J., Lehar, J., Kryukov, G.V., Sonkin, D., et al. (2012). The Cancer Cell Line Encyclopedia enables predictive modelling of anticancer drug sensitivity. Nature 483, 603–607.

Basturk, O., Tang, L., Hruban, R.H., Adsay, V., Yang, Z., Krasinskas, A.M., Vakiani, E., La Rosa, S., Jang, K.T., Frankel, W.L., et al. (2014). Poorly differentiated neuroendocrine carcinomas of the pancreas: a clinicopathologic analysis of 44 cases. Am J Surg Pathol 38, 437–447.

Belle, N.M., Ji, Y., Herbine, K., Wei, Y., Park, J., Zullo, K., Hung, L.Y., Srivatsa, S., Young, T., Oniskey, T., et al. (2019). TFF3 interacts with LINGO2 to regulate EGFR activation for protection against colitis and gastrointestinal helminths. Nat Commun 10, 4408.

Bosman, F.T., World Health Organization., and International Agency for Research on Cancer. (2010). WHO classification of tumours of the digestive system, 4th edn (Lyon: International Agency for Research on Cancer).

Castillo-Azofeifa, D., Fazio, E.N., Nattiv, R., Good, H.J., Wald, T., Pest, M.A., de Sauvage, F.J., Klein, O.D., and Asfaha, S. (2019). Atoh1(+) secretory progenitors possess renewal capacity independent of Lgr5(+) cells during colonic regeneration. EMBO J 38.

Cheng, Y., Liao, S., Xu, G., Hu, J., Guo, D., Du, F., Contreras, A., Cai, K.Q., Peri, S., Wang, Y., et al. (2020). NeuroD1 Dictates Tumor Cell Differentiation in Medulloblastoma. Cell Rep 31, 107782.

De Roock, W., Claes, B., Bernasconi, D., De Schutter, J., Biesmans, B., Fountzilas, G., Kalogeras, K.T., Kotoula, V., Papamichael, D., Laurent-Puig, P., et al. (2010). Effects of KRAS, BRAF, NRAS, and PIK3CA mutations on the efficacy of cetuximab plus chemotherapy in chemotherapy-refractory metastatic colorectal cancer: a retrospective consortium analysis. Lancet Oncol 11, 753–762.

Fiegl, H., Jones, A., Hauser-Kronberger, C., Hutarew, G., Reitsamer, R., Jones, R.L., Dowsett, M., Mueller-Holzner, E., Windbichler, G., Daxenbichler, G., et al. (2008). Methylated NEUROD1 promoter is a marker for chemosensitivity in breast cancer. Clin Cancer Res 14, 3494–3502.

Fleming, M., Ravula, S., Tatishchev, S.F., and Wang, H.L. (2012). Colorectal carcinoma: Pathologic aspects. J Gastrointest Oncol 3, 153–173.

Gehart, H., van Es, J.H., Hamer, K., Beumer, J., Kretzschmar, K., Dekkers, J.F., Rios, A., and Clevers, H. (2019). Identification of Enteroendocrine Regulators by Real-Time Single-Cell Differentiation Mapping. Cell 176, 1158–1173 e1116.

Harada, A., Teng, J., Takei, Y., Oguchi, K., and Hirokawa, N. (2002). MAP2 is required for dendrite elongation, PKA anchoring in dendrites, and proper PKA signal transduction. J Cell Biol 158, 541–549.

Henry, C., Close, A.F., and Buteau, J. (2014). A critical role for the neural zinc factor ST18 in pancreatic beta-cell apoptosis. J Biol Chem 289, 8413–8419.

Hsu, H.C., Liu, Y.S., Tseng, K.C., Yang, T.S., Yeh, C.Y., You, J.F., Hung, H.Y., Chen, S.J., and Chen, H.C. (2015). CBB1003, a lysine-specific demethylase 1 inhibitor, suppresses colorectal cancer cells growth through down-regulation of leucine-rich repeat-containing G-protein-coupled receptor 5 expression. J Cancer Res Clin Oncol 141, 11–21.

Huet, C., Sahuquillo-Merino, C., Coudrier, E., and Louvard, D. (1987). Absorptive and mucus-secreting subclones isolated from a multipotent intestinal cell line (HT-29) provide new models for cell polarity and terminal differentiation. J Cell Biol 105, 345–357.

Hugen, N., van Beek, J.J., de Wilt, J.H., and Nagtegaal, I.D. (2014). Insight into mucinous colorectal carcinoma: clues from etiology. Ann Surg Oncol 21, 2963–2970.

Inoue, Y., Horie, H., Homma, Y., Sadatomo, A., Tahara, M., Koinuma, K., Yamaguchi, H., Mimura, T., Kihara, A., Lefor, A.K., et al. (2020). Goblet cell carcinoid of the rectum: a case report. Surg Case Rep 6, 174.

Jassal, B., Matthews, L., Viteri, G., Gong, C., Lorente, P., Fabregat, A., Sidiropoulos, K., Cook, J., Gillespie, M., Haw, R., et al. (2020). The reactome pathway knowledgebase. Nucleic Acids Res 48, D498–D503.

Jensen, J., Pedersen, E.E., Galante, P., Hald, J., Heller, R.S., Ishibashi, M., Kageyama, R., Guillemot, F., Serup, P., and Madsen, O.D. (2000). Control of endodermal endocrine development by Hes-1. Nat Genet 24, 36–44.

Laurent, B., Ruitu, L., Murn, J., Hempel, K., Ferrao, R., Xiang, Y., Liu, S., Garcia, B.A., Wu, H., Wu, F., et al. (2015). A specific LSD1/KDM1A isoform regulates neuronal differentiation through H3K9 demethylation. Mol Cell 57, 957–970.

Lin, J.Z., and Farmer, S.R. (2016). LSD1-a pivotal epigenetic regulator of brown and beige fat differentiation and homeostasis. Genes Dev 30, 1793–1795.

Maiques-Diaz, A., Spencer, G.J., Lynch, J.T., Ciceri, F., Williams, E.L., Amaral, F.M.R., Wiseman, D.H., Harris, W.J., Li, Y., Sahoo, S., et al. (2018). Enhancer Activation by Pharmacologic Displacement of LSD1 from GFI1 Induces Differentiation in Acute Myeloid Leukemia. Cell Rep 22, 3641–3659.

Mao, M., Tian, F., Mariadason, J.M., Tsao, C.C., Lemos, R., Jr., Dayyani, F., Gopal, Y.N., Jiang, Z.Q., Wistuba, II, Tang, X.M., et al. (2013). Resistance to BRAF inhibition in BRAF-mutant colon cancer can be overcome with PI3K inhibition or demethylating agents. Clin Cancer Res 19, 657–667.

Margonis, G.A., Buettner, S., Andreatos, N., Kim, Y., Wagner, D., Sasaki, K., Beer, A., Schwarz, C., Loes, I.M., Smolle, M., et al. (2018). Association of BRAF Mutations With Survival and Recurrence in Surgically Treated Patients With Metastatic Colorectal Liver Cancer. JAMA Surg 153, e180996.

McGory, M.L., Maggard, M.A., Kang, H., O’Connell, J.B., and Ko, C.Y. (2005). Malignancies of the appendix: beyond case series reports. Dis Colon Rectum 48, 2264–2271.

Miller, S.A., Policastro, R.A., Savant, S.S., Sriramkumar, S., Ding, N., Lu, X., Mohammad, H.P., Cao, S., Kalin, J.H., Cole, P.A., et al. (2020). Lysine-Specific Demethylase 1 Mediates AKT Activity and Promotes Epithelial-to-Mesenchymal Transition in PIK3CA-Mutant Colorectal Cancer. Mol Cancer Res 18, 264–277.

Modlin, I.M., and Sandor, A. (1997). An analysis of 8305 cases of carcinoid tumors. Cancer 79, 813–829.

Moertel, C.G., Dockerty, M.B., and Judd, E.S. (1968). Carcinoid tumors of the vermiform appendix. Cancer 21, 270–278.

Mohammad, H.P., Smitheman, K.N., Kamat, C.D., Soong, D., Federowicz, K.E., Van Aller, G.S., Schneck, J.L., Carson, J.D., Liu, Y., Butticello, M., et al. (2015). A DNA Hypomethylation Signature Predicts Antitumor Activity of LSD1 Inhibitors in SCLC. Cancer Cell 28, 57–69.

Peignon, G., Durand, A., Cacheux, W., Ayrault, O., Terris, B., Laurent-Puig, P., Shroyer, N.F., Van Seuningen, I., Honjo, T., Perret, C., et al. (2011). Complex interplay between beta-catenin signalling and Notch effectors in intestinal tumorigenesis. Gut 60, 166–176.

Ray, S.K., Li, H.J., Metzger, E., Schule, R., and Leiter, A.B. (2014). CtBP and associated LSD1 are required for transcriptional activation by NeuroD1 in gastrointestinal endocrine cells. Mol Cell Biol 34, 2308–2317.

Reischmann, N., Andrieux, G., Griffin, R., Reinheckel, T., Boerries, M., and Brummer, T. (2020). BRAF(V600E) drives dedifferentiation in small intestinal and colonic organoids and cooperates with mutant p53 and Apc loss in transformation. Oncogene 39, 6053–6070.

Sakamoto, N., Feng, Y., Stolfi, C., Kurosu, Y., Green, M., Lin, J., Green, M.E., Sentani, K., Yasui, W., McMahon, M., et al. (2017). BRAF(V600E) cooperates with CDX2 inactivation to promote serrated colorectal tumorigenesis. Elife 6.

Schlesinger, Y., Straussman, R., Keshet, I., Farkash, S., Hecht, M., Zimmerman, J., Eden, E., Yakhini, Z., Ben-Shushan, E., Reubinoff, B.E., et al. (2007). Polycomb-mediated methylation on Lys27 of histone H3 pre-marks genes for de novo methylation in cancer. Nat Genet 39, 232–236.

Schneider, J.W., Gao, Z., Li, S., Farooqi, M., Tang, T.S., Bezprozvanny, I., Frantz, D.E., and Hsieh, J. (2008). Small-molecule activation of neuronal cell fate. Nat Chem Biol 4, 408–410.

Schneider, N.I., and Langner, C. (2014). Prognostic stratification of colorectal cancer patients: current perspectives. Cancer Manag Res 6, 291–300.

Schonhoff, S.E., Giel-Moloney, M., and Leiter, A.B. (2004). Neurogenin 3-expressing progenitor cells in the gastrointestinal tract differentiate into both endocrine and non-endocrine cell types. Dev Biol 270, 443–454.

Simmini, S., Bialecka, M., Huch, M., Kester, L., van de Wetering, M., Sato, T., Beck, F., van Oudenaarden, A., Clevers, H., and Deschamps, J. (2014). Transformation of intestinal stem cells into gastric stem cells on loss of transcription factor Cdx2. Nat Commun 5, 5728.

Takagi, S., Ishikawa, Y., Mizutani, A., Iwasaki, S., Matsumoto, S., Kamada, Y., Nomura, T., and Nakamura, K. (2017). LSD1 Inhibitor T-3775440 Inhibits SCLC Cell Proliferation by Disrupting LSD1 Interactions with SNAG Domain Proteins INSM1 and GFI1B. Cancer Res 77, 4652–4662.

Taupin, D.R., Kinoshita, K., and Podolsky, D.K. (2000). Intestinal trefoil factor confers colonic epithelial resistance to apoptosis. Proc Natl Acad Sci U S A 97, 799–804.

Tomic, G., Morrissey, E., Kozar, S., Ben-Moshe, S., Hoyle, A., Azzarelli, R., Kemp, R., Chilamakuri, C.S.R., Itzkovitz, S., Philpott, A., et al. (2018). Phospho-regulation of ATOH1 Is Required for Plasticity of Secretory Progenitors and Tissue Regeneration. Cell Stem Cell 23, 436–443 e437.

Tosic, M., Allen, A., Willmann, D., Lepper, C., Kim, J., Duteil, D., and Schule, R. (2018). Lsd1 regulates skeletal muscle regeneration and directs the fate of satellite cells. Nat Commun 9, 366.

Toyota, M., Ahuja, N., Ohe-Toyota, M., Herman, J.G., Baylin, S.B., and Issa, J.P. (1999). CpG island methylator phenotype in colorectal cancer. Proc Natl Acad Sci U S A 96, 8681–8686.

Uhlitz, F., Bischoff, P., Sieber, A., Obermayer, B., Blanc, E., Lüthen, M., Sawitzki, B., Kamphues, C., Beule, D., Sers, C., et al. (2020). A census of cell types and paracrine interactions in colorectal cancer. bioRxiv.

van Eeden, S., Offerhaus, G.J., Hart, A.A., Boerrigter, L., Nederlof, P.M., Porter, E., and van Velthuysen, M.L. (2007). Goblet cell carcinoid of the appendix: a specific type of carcinoma. Histopathology 51, 763–773.

Vogelstein, B., Papadopoulos, N., Velculescu, V.E., Zhou, S., Diaz, L.A., Jr., and Kinzler, K.W. (2013). Cancer genome landscapes. Science 339, 1546–1558.

Weisenberger, D.J., Siegmund, K.D., Campan, M., Young, J., Long, T.I., Faasse, M.A., Kang, G.H., Widschwendter, M., Weener, D., Buchanan, D., et al. (2006). CpG island methylator phenotype underlies sporadic microsatellite instability and is tightly associated with BRAF mutation in colorectal cancer. Nat Genet 38, 787–793.

Yaeger, R., Cercek, A., Chou, J.F., Sylvester, B.E., Kemeny, N.E., Hechtman, J.F., Ladanyi, M., Rosen, N., Weiser, M.R., Capanu, M., et al. (2014). BRAF mutation predicts for poor outcomes after metastasectomy in patients with metastatic colorectal cancer. Cancer 120, 2316–2324.

Yaeger, R., Cercek, A., O’Reilly, E.M., Reidy, D.L., Kemeny, N., Wolinsky, T., Capanu, M., Gollub, M.J., Rosen, N., Berger, M.F., et al. (2015). Pilot trial of combined BRAF and EGFR inhibition in BRAF-mutant metastatic colorectal cancer patients. Clin Cancer Res 21, 1313–1320.

Yang, Q., Bermingham, N.A., Finegold, M.J., and Zoghbi, H.Y. (2001). Requirement of Math1 for secretory cell lineage commitment in the mouse intestine. Science 294, 2155–2158.

Zwiggelaar, R.T., Lindholm, H.T., Fosslie, M., Terndrup Pedersen, M., Ohta, Y., Diez-Sanchez, A., Martin-Alonso, M., Ostrop, J., Matano, M., Parmar, N., et al. (2020). LSD1 represses a neonatal/reparative gene program in adult intestinal epithelium. Sci Adv 6.

