## Supplemental Materials and Methods for "LSD1 promotes secretory cell specification to drive *BRAF* mutant colorectal cancer"

**Supplemental Methods and Materials**

**TCGA Analyses**

We acquired TCGA COAD datasets from Xena Browser. 383 patient tumors met our specifications at the beginning of the analysis, including mutation and expression data. The scripts used for TCGA data analysis can be found on github (https://github.com/rpolicastro/Miller_Policastro_2020_TCGA). All multiple comparisons for TCGA analysis were corrected using Benjamini-Hochberg unless otherwise stated. Counts were squared and then TMM normalized using edgeR(McCarthy et al., 2012). For the quintile comparison plot in Fig 1B and Supplemental Fig 1D, the selected genes were first binned into quintiles based on their relative expression across samples, and then samples were separated based on putative driver mutation. For each displayed gene pair and driver mutation, the number of samples overlapping each quintile bin combination was plotted. The associated p-values for Supplemental Fig 1B & E were calculated based on a permutation monte-carlo simulation. The quintile labels were randomly shuffled for each gene, and then the simulated quintile overlaps between each gene pair for each driver mutation were recomputed for a total of 10,000 iterations. The p-values were determined based on the number of simulated counts that were equal to or greater than the observed count plus one over the number of iterations plus one (Phipson and Smyth, 2010) and were then corrected for multiple comparisons using FDR. For the correlation plots in Supplemental Fig 1A & C, the TMM normalized counts were $Log_2 + 1$ transformed and then the Pearson correlation coefficients and p-values were calculated using the R function *cor.test*. The regression line and confidence interval were added to the plot using the *geom_smooth* layer from the R plotting library ggplot2.

The euler plot in Fig 1C was generated using the eulerr R library(Larsson, 2020). Samples were first categorized based on both the mutation status of *BRAF* and *KRAS*, and then by having expression in the top 2 quintiles for *CHGA* and *SST* ($BRAF^{EC}$ or $KRAS^{EC}$) or *NEUROG3* and *INSM1* ($BRAF^{EP}$ or $KRAS^{EP}$). For the boxplot in Fig 1D, the categories from the euler plot were used in addition to matched normal tissue and all other uncategorized tumors. For each gene, pairwise Wilcoxon rank-sum tests were calculated between the TMM normalized expression values of the gene in all sample subtypes. For display in the plot, the gene expression values were Log2 + 1 transformed. For the jitter plot in Supplemental Fig 1F, the expression values of *MUC2* and *ATOH1* were plotted based on subtype, and points were colored by tumor diagnosis. Expression values between subtypes were compared using pairwise Wilcoxon rank-sum tests, and differences in diagnosis counts between subtypes were computed using pairwise Fisher exact tests. The TCGA DNA methylation array data for Fig 3A and Supplemental Fig 5 was merged into the tumor subtype categories, and gene tracks of average methylation levels for each subtype were created with the R bioconductor library Gviz(Hahne and Ivanek, 2016).

**Single-Cell Sequencing**

HT29 and H508 samples were sequenced to a minimum of 3000 cells with approximately 30,000 sequencing reads per cell to allow capture of rare and transient cell types, as well as the expression profiles of approximately 5,000 genes per cell on average. To compare the cell lines against the normal human large intestine, we also searched the literature for high-quality normal colon scRNA-seq datasets. The quality of datasets was determined by comparing the number of detected genes per cell versus the total percentage of mitochondrial transcript content of the cell (Supplemental Figure 3). Using these criteria, we selected one normal human colon dataset which contained approximately 1,600 cells passing quality control, with an average of 3,000 genes detected and 18,000 sequencing reads per cell.

All scripts used for scRNA-seq analysis are available on github (https://github.com/rpolicastro/Miller_Policastro_2020_scRNAseq). 10X Genomics Cell Ranger (v3.1.0) with default settings was used for quality control, alignment, and feature counting of demultiplexed reads using the GRCh38 (v3.0.0) genome assembly provided by 10X Genomics. For RNA velocity, the BAM files from Cell Ranger were processed with velocyto (v0.17.17)(La Manno et al., 2018) to obtain the spliced, unspliced, and ambiguous feature UMI counts. The matrices from Cell Ranger and velocyto were then read into Seurat (v3.1.5) (Butler et al., 2018) for further processing. For the Colon sample, cells were filtered with less than 1000 genes detected or greater than 30% mitochondrial RNA content, and for the HT29 and H508 samples, cells were filtered with less than 2500 genes detected or more than 25% mitochondrial RNA content. The cell-cycle stage was annotated using the Seurat function *CellCycleScoring* using the built-in genes in the variable *cc.genes*. UMI counts were then normalized using SCTransform(Hafemeister and Satija, 2019) with default settings, and then integrated(Stuart et al., 2019) using the HT29 and H508 EV samples as references. Clusters were determined in Seurat using the Leiden algorithm(Traag et al., 2019) and refined after marker gene exploration (Wilcoxon rank-sum test; FDR < 0.05 and Log Fold Change > 0.4), and UMAP(McInnes, 2020) (Becht et al., 2018) dimension reduction was used to display the cells in two dimensions. The processed scRNA-seq is available for browsing at http://185.215.224.30:3838/miller/miller_policastro_2020/, which was also used to generate all UMAP plots. All plots with gene counts use the normalized and Log2 transformed UMI counts from Seurat.

The code to find the proportional difference in cell populations between two samples is available as an R library on github (https://github.com/rpolicastro/scProportionTest/releases/tag/v1.0.0). The significance of the difference was calculated by permutation testing. Briefly, the cell identities were randomly shuffled between the two samples, and the Log2 proportional difference between the cell counts in each population was found. This process was repeated 10000 times and the p-values were calculated by the number of proportional differences that were as or more extreme than observed plus one, over the iteration number plus one(Phipson and Smyth, 2010). The resulting p-values were then FDR corrected for multiple comparisons. Bootstrap sampling was used to generate 95% confidence intervals for plotting. Cell identities for each sample were sampled with replacement, and then the Log2 fold difference between cell counts in each population were obtained. This was repeated 10000 times, and the confidence interval was the 0.025 to 0.975 quantile range of simulated Log2 proportional differences in cell counts.

After preprocessing the Velocyto output in Seurat, the corrected counts, clusters, and UMAP embeddings were loaded into scVelo (v0.2.1)(Bergen et al., 2020), the ratio of spliced to unspliced reads per cluster was found, and cell velocities were computed. All functions were run with default settings unless otherwise stated. The *scvelo.pp.filter_and_normalize* argument 'n_top_genes' was set to 3000, and the 'n_npcs' and 'n_neighbors' arguments of *scvelo.pp.momentum* were both set to 30. The velocity streamlines and cell arrows were made with the *scvelo.pl.velocity_embedding_stream* and *scvelo.pl.velocity_embedding functions*. The top velocity genes per cluster were discovered using *scvelo.tl.rank_velocity_genes*, and plotted using *scvelo.pl.velocity*. *scvelo.tl.velocity_confidence* generated the velocity confidence and length values, and the results were plotted using *scvelo.pl.scatter*. Cell transition states were traced and plotted using scvelo.utils.get_cell_transition, *scvelo.pl.umap*, and *scvelo.pl.scatter*. The PAGA velocity graph was generated with *scvelo.pl.paga(Wolf et al., 2019)*. Finally, the end state cell and cluster plots were generated using the CellRank (rc v1.8.0) functions *cellrank.pl.terminal_states* and *g.plot_metastable_states* with default kernel parameters.

**Western Blot Antibodies**

LSD1 (CST: 2139) (Western blot), AKT (CST: 4691) (Western blot), phospho-Ser473-AKT (CST: 4060) (Western blot), (Western blot), B-actin (CST: 4970) (Western blot), phospho-TSC2 (Ser1387) (CST: 23402) (Western blot), Snail (CST: 3879) (Western blot), secondary Alexa conjugate (CST: mouse 8890 and rabbit 8889) (Western blot), Alexa flour anti-rabbit 594, Alexa flour anti-mouse 488, (CST: rabbit 8889 and mouse 44008)

**qPCR Primer sequences listed below:**

*LSD1*, forward, GGTGAGCTCTTCCTCTTCTGG;

*LSD1*, reverse, TCGGCCAACAATCACATCGT;

*ATOH1*, forward, AGAGAGCATCCCGTCTACCC;

*ATOH1*, reverse, GCTCCGGGGAATGTAGCAAA;

*NEUROG3*, forward, CGGTAGAAAGGATGACGCCT;

*NEUROG3*, reverse, GGTCACTTCGTCTTCCGAGG;

*NEUROD1*, forward, GACACGAGGAATTCGCCCA;

*NEUROD1*, reverse, CCCACTCTCGCTGTACGATT;

*RHOA*, forward, CGTTAGTCCACGGTCTGGTC;

*RHOA*, reverse, ACCAGTTTCTTCCGGATGGC;

*B-ACTIN*, forward, GAAGCCGGCCTTGCACAT;

*B-ACTIN*, reverse, AGCACAGAGCCTCGCCTTT;

*TFF3*, forward, GGAGTGCCTTGGTGTTTCAAG;

*TFF3*, reverse, AAAGCTGAGATGAACAGTGCCT;

*SPDEF*, forward, CTCAGCTGCCCACACCTCTT;

*SPDEF*, reverse, GGGATACGCTGCTCAGACC;

*HES1*, forward, AAAAATTCCTCGTCCCCGGT;

*HES1*, reverse, GGCTTTGATGACTTTCTGTGCT.

**qMSP primer sequences listed below:**

*NEUROD1,* LM, CGGTTTAGTTAAGAGTTTGGAATTAC

*NEUROD1,* RM, CTACGAATAAAAACAAATCCGC

*NEUROD1,* LU, TTGGTTTAGTTAAGAGTTTGGAATTATG

*NEUROD1,* RU, TCTCCTACAAATAAAAACAAATCCAC

| | Cell Lines | | | |
|---|---|---|---|---|
| | HT29 | NCI-H508 | LS174T | SW480 |
| Cancer Subtype | Adeno-carcinoma (primary) | Adeno-carcinoma (ascites met.) | Adeno-carcinoma (primary) | Adeno-carcinoma (primary) |
| *BRAF* | V600E | G596R | WT | WT |
| *KRAS* | WT | WT | G12D | G12V |
| *APC* | E853* | WT | WT | Q1338* |
| *PIK3CA* | P449T | E545K | H1047R | WT |
| *TP53* | R273H | R273H | WT | R273H P309S |
| *SMAD4* | Q311* | WT | WT | WT |

**Supplemental Table 1: Common mutations in cell lines used for this study (mutations obtained from the CCLE(Barretina et al., 2012)).**

Excel Worksheet

**Supplemental Table 2: Statistical analysis of TCGA COAD comparing Normal, *BRAF* mutant, *KRAS* mutant, and all other tumors.**

Excel Worksheet

**Supplemental Table 3: Statistical analysis of *ATOH1* and *MUC2* expression in TCGA COAD comparing Normal, *BRAF* mutant, *KRAS* mutant, and all other tumors.**

Excel Worksheet

**Supplemental Table 4: Statistical analysis of pathological subtype enrichment in TCGA COAD *BRAF* mutant, *KRAS* mutant, and all other tumor groups.**

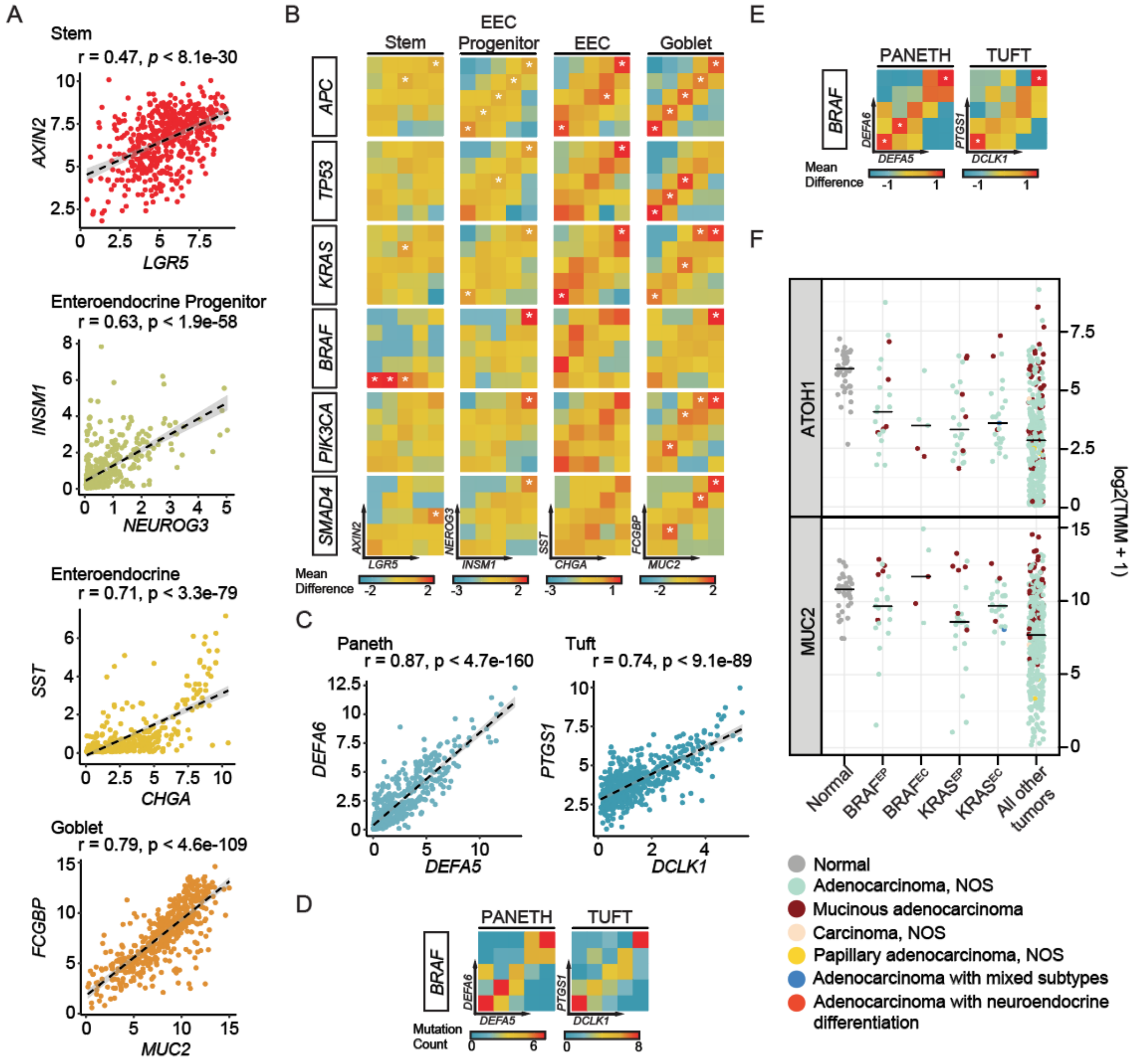


**Supplemental Figure 1: TCGA validation of cell-type markers and analysis of pathological and molecular features.**

A, Correlation plots for cell-type-specific markers in TCGA COAD dataset, where r = Pearson's correlation coefficient. B, Coexpression of cell-type marker genes plotted by mutation type and visualized through quantile plots of mean difference in quantile enrichment. Stars depict quantiles with enrichment FDR < 0.05. C, Correlation plots for additional cell-type-specific markers in TCGA COAD dataset, where r = Pearson's correlation coefficient. D, Coexpression of cell-type marker genes plotted by mutation type and visualized through quantile plots of absolute counts. E, Coexpression of cell-type marker genes plotted by mutation type and visualized through quantile plots of mean difference in quantile enrichment. Stars depict quantiles with enrichment FDR < 0.05. F, Dotplot with indicated means, showing expression of *ATOH1* or *MUC2* in normal tissue (Normal), BRAF$^{EP}$, BRAF$^{EC}$, KRAS$^{EP}$, KRAS$^{EC,}$ and all other

COAD dataset tumors (All other tumors). Individual samples are colored based on pathological classification.

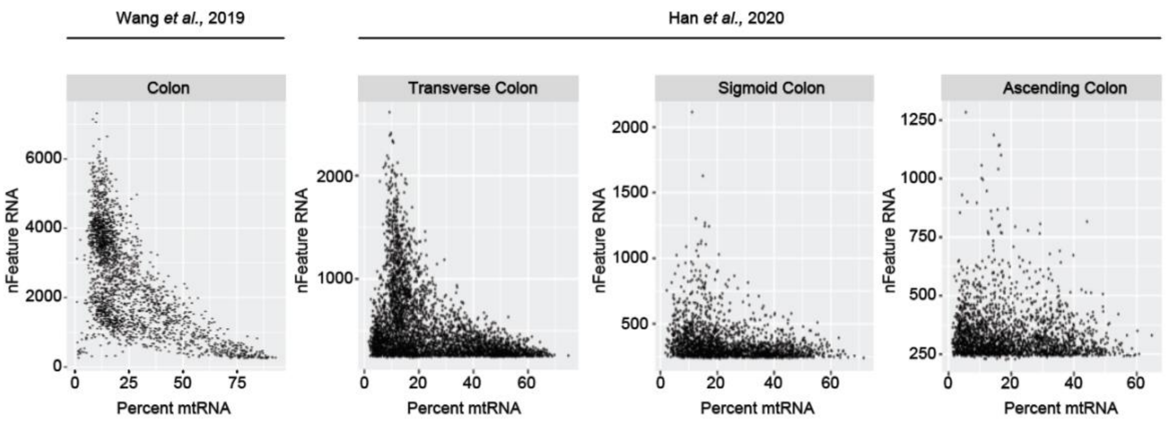


**Supplemental Figure 2: Comparison of sequencing quality among published human colon scRNA-seq datasets.**

Scatterplots depicting the number of detected genes (nFeature RNA) versus the percentage of reads derived from mitochondrial genes (Percent mtRNA) from published 10X Chromium scRNA-seq(Wang et al., 2020) and Microwell-seq(Han et al., 2020) datasets.

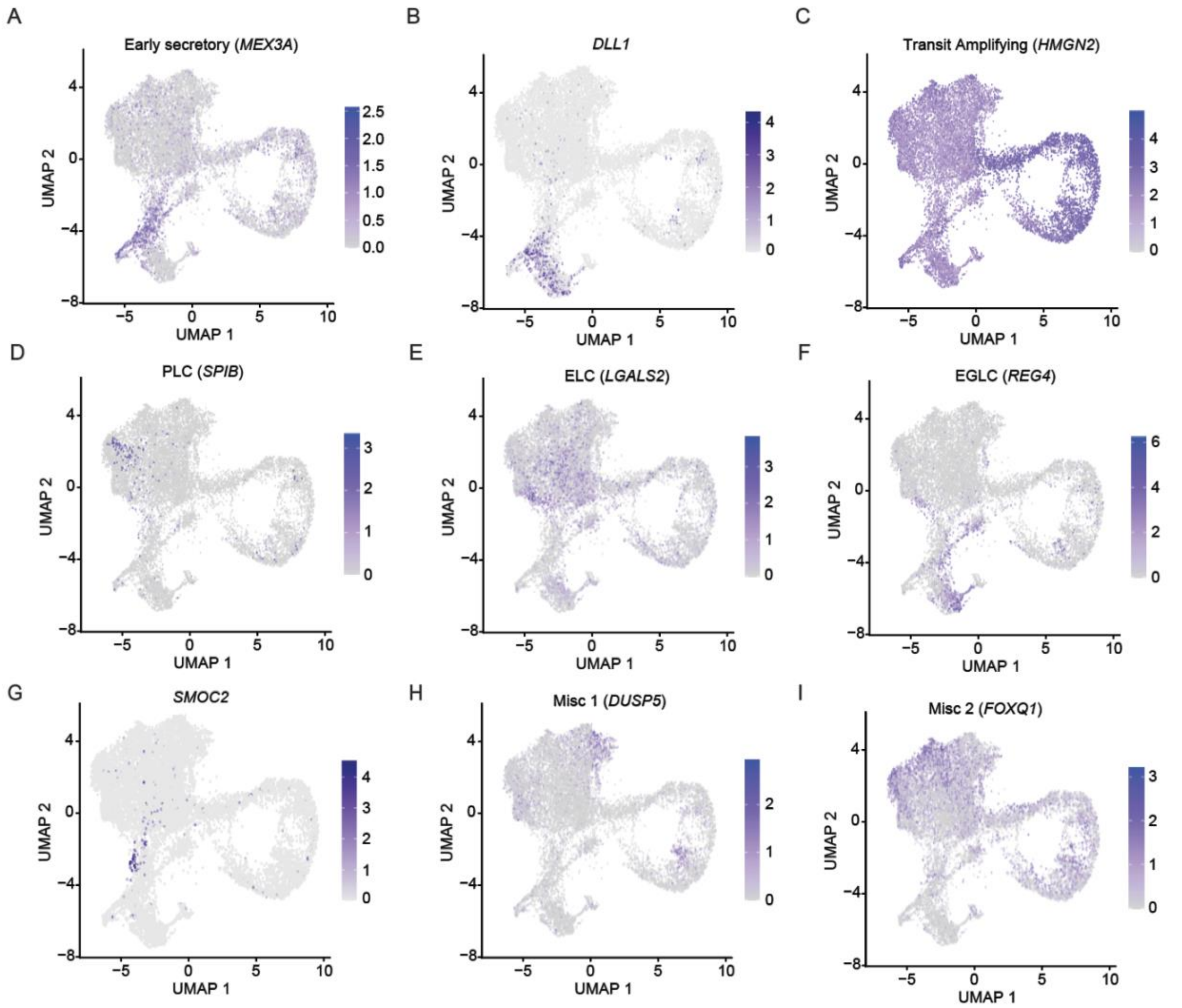


**Supplemental Figure 3: Cell line and colon subpopulations express canonical cell-type-specific gene markers.**

UMAP dot plots of normalized expression values of marker genes representative of the (A) EGLC, (B) Early secretory, (C) PLC, (D) ELC, (E) Misc 1, (F) Misc 2, (G) Transit Amplifying cells in the colon, as well as (H, I) select markers in the colon, HT29 EV and H508 EV samples.

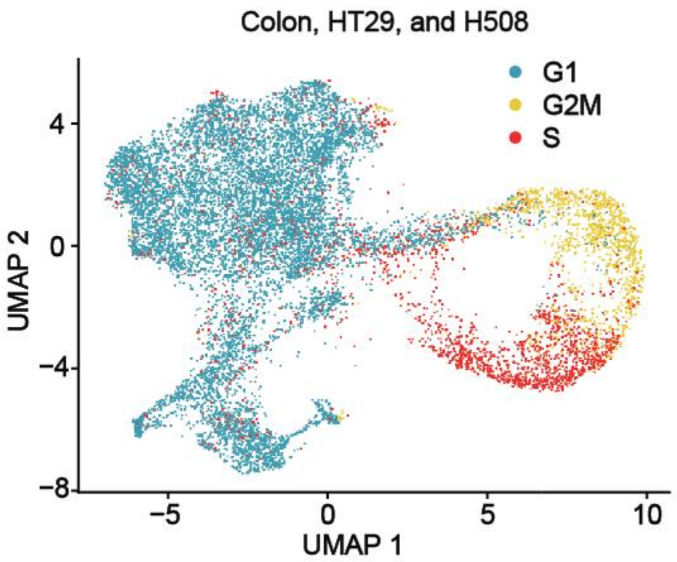


**Supplemental Figure 4: Cell-cycle analysis of scRNA-seq datasets.**

UMAP dot plot detailing predicted cell cycle phase of normal human colon, HT29 EV, and H508 EV scRNA-seq samples.

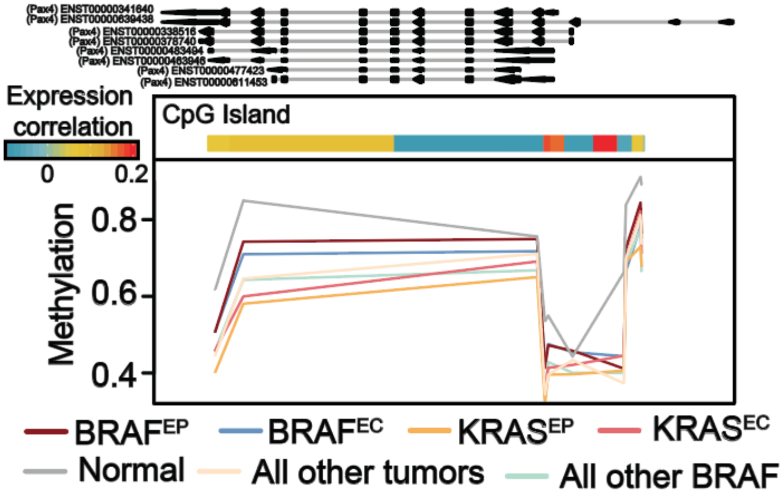


**Supplemental Figure 5: *NEUROD1* CpG island methylation in *BRAF* mutant CRC abrogates *NEUROD1* expression.**

Mean percent methylation from TCGA COAD Illumina human methylation 450 array probes across the transcription start site of *PAX4.* The expression correlation heatmap depicts Spearman’s correlation between *NEUROD1* mRNA expression and 450K array methylation.

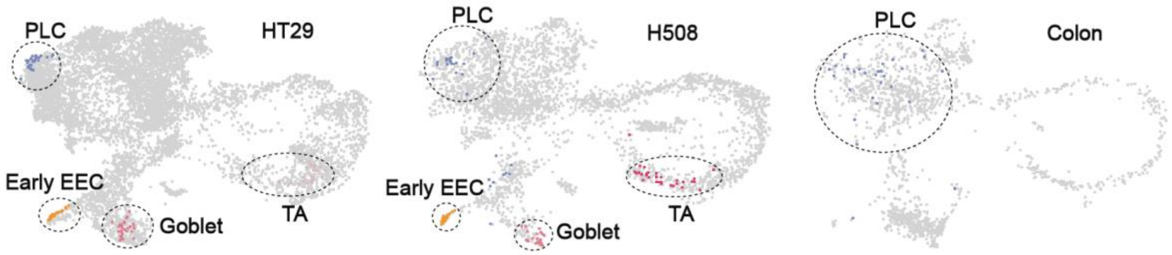


**Supplemental Figure 6: CellRank predicted end-states in each scRNA-seq sample.**

CellRank prediction of cell differentiation end-states in HT29, H508, and colon scRNA-seq datasets. Dotted circles emphasize clusters predicted to have end-state cells.

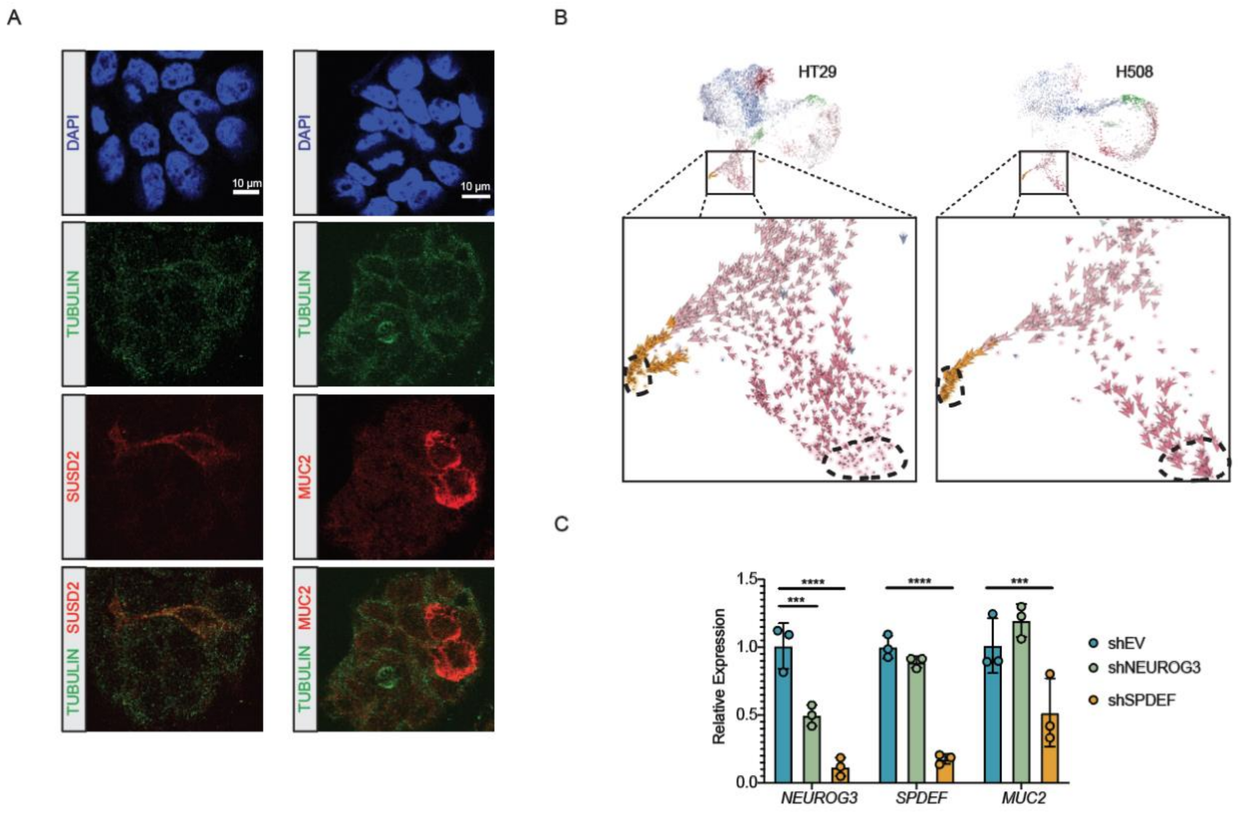


**Supplemental Figure 7: Molecular validation of RNA velocity predictions.**

A, Representative immunofluorescence images of HT29 cells depicting SUSD2 (EEC progenitor marker) or MUC2 (goblet marker) signal. B, Velocity arrows for individual cells in the early secretory, goblet, and early EEC cell clusters, with a dotted circle indicating predicted terminal cell states in each subpopulation. C, RT-qPCR analysis of *NEUROG3*, *SPDEF*, and *MUC2* RNA expression level in EV control or NEUROG3 KD, SPDEF KD cells. Significance determined by two-way ANOVA with pairwise Dunnett’s multiple comparisons testing (***, P<0.001; ****, P<0.0001).

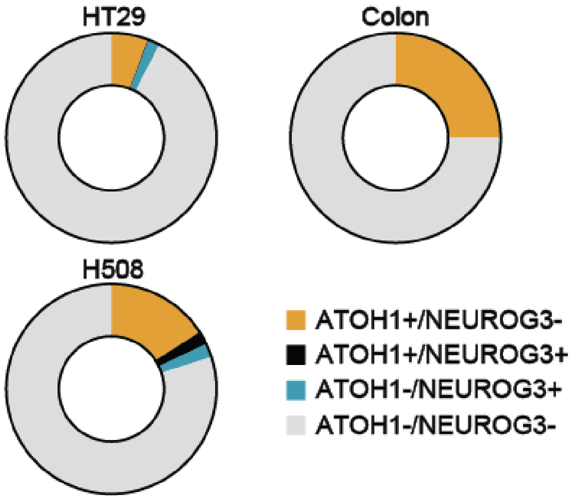


**Supplemental Figure 8. Analysis of secretory progenitor proportions in scRNA-seq datasets.**

Donut chart depicting the proportions of cells expressing various combinations of *ATOH1* and *NEUROG3* in HT29, H508, and colon scRNA-seq datasets.

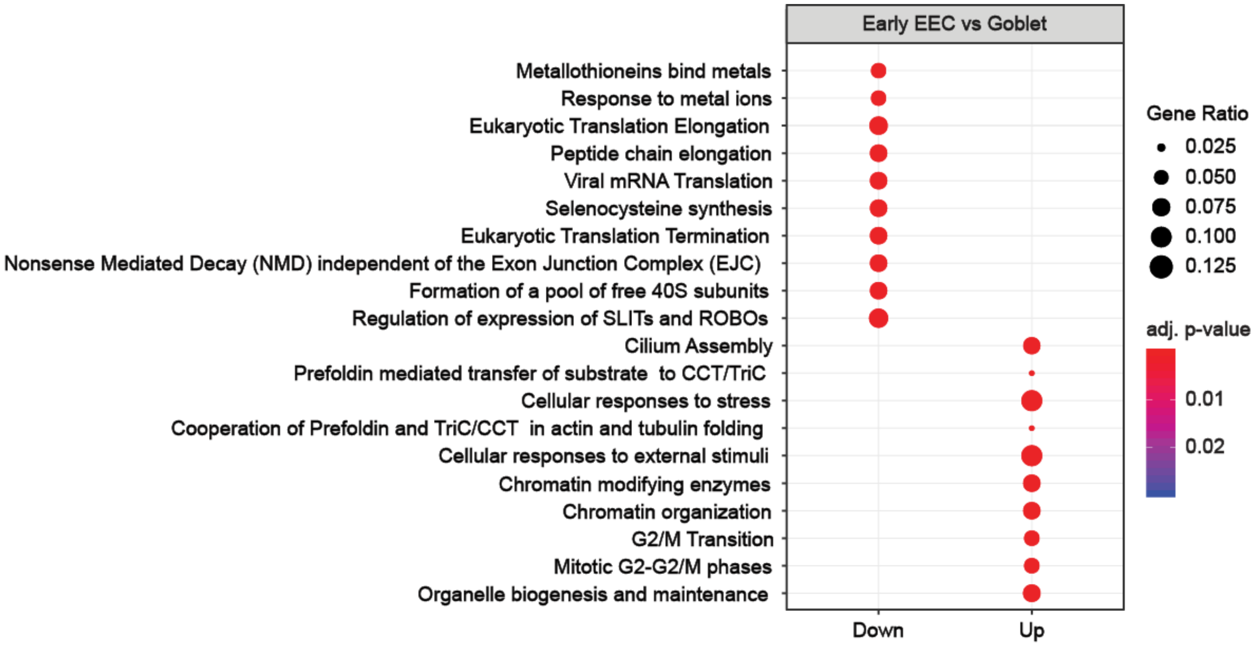


**Supplemental Figure 9: Early EEC and goblet cell expression profiles are enriched for unique functional pathways.**

Pathway analysis of differentially expressed genes between early EEC and goblet subpopulations. The size of the circle denotes the proportion of term genes represented in the dataset, and color represents the degree of significance by adj. p-value.

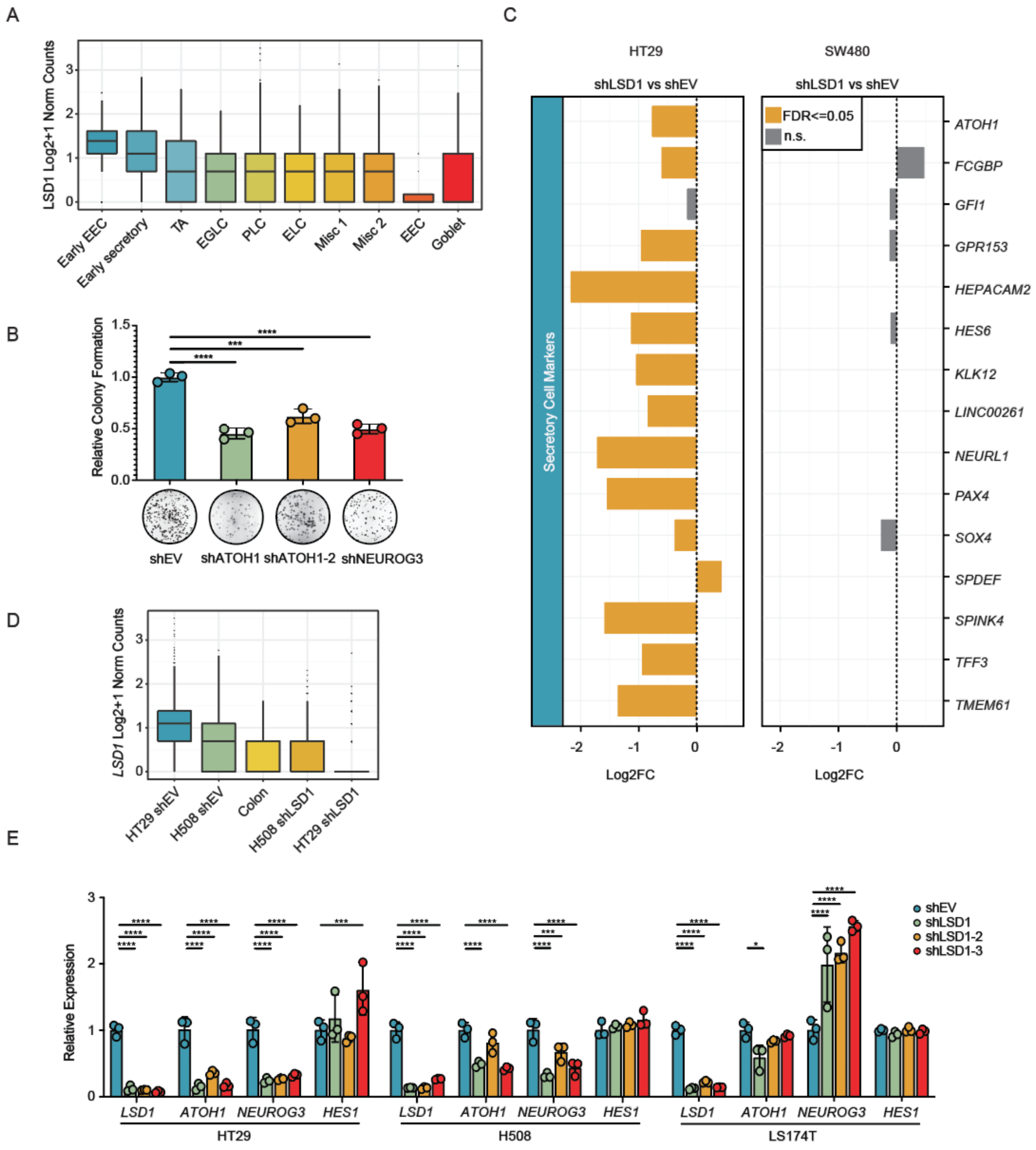


**Supplemental Figure 10. *LSD1* levels in single-cell RNA-sequencing datasets and analysis of LSD1 KD in bulk RNA-seq.**

A, Expression of *LSD1* in subpopulations of the integrated colon, HT29, and H508 datasets. B, Assay of clonogenic growth in EV, ATOH1 KD, ATOH1-2 KD, and NEUROG3 KD HT29 cells. C,

Barplot depicts Log2FC in gene expression between LSD1 KD and EV in RNA-seq datasets from HT29 or SW480 cell lines. Orange bars depict significant changes in gene expression (FDR <= 0.05) and grey bars are not significant. D, Bulk *LSD1* expression in all single-cell RNA-seq samples. E, RT-qPCR analysis of *LSD1*, *ATOH1*, *NEUROG3*, and *HES1* RNA expression level in EV control or three independent LSD1 KDs. Significance determined by (B, E) one-way ANOVA with pairwise Dunnett's multiple comparisons testing (*, P<0.05; ***, P<0.001; ****, P<0.0001).

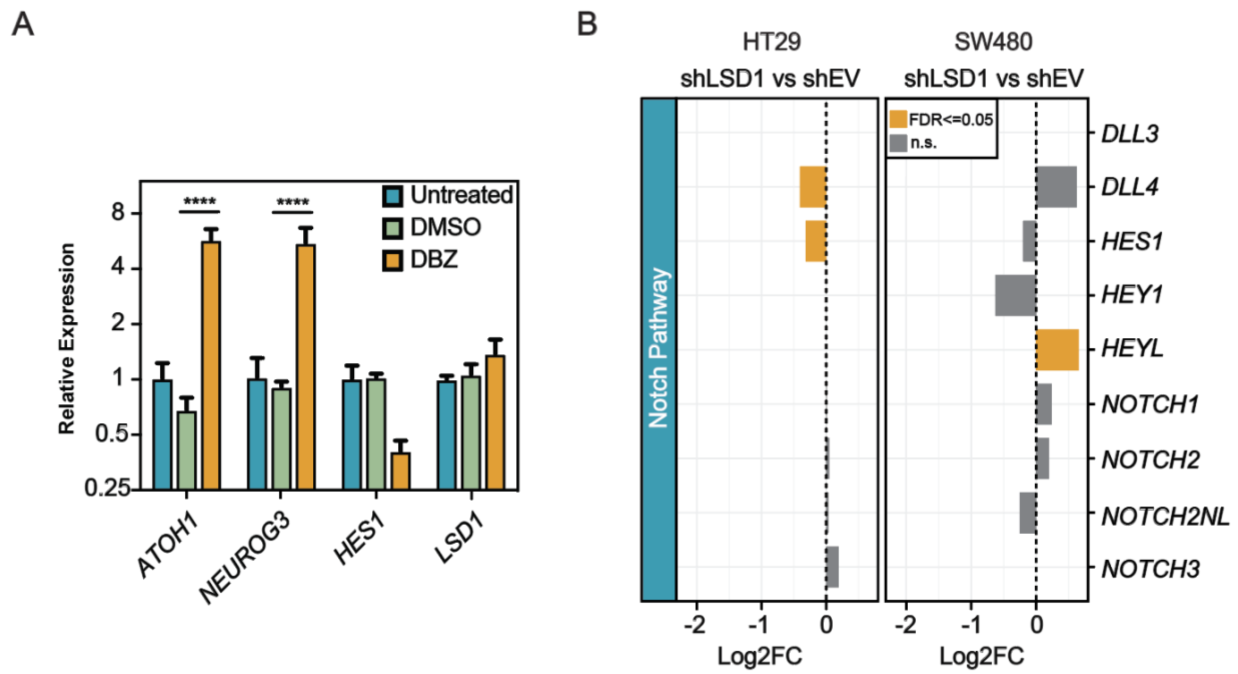


**Supplemental Figure 11: Examination of the Notch pathway in LSD1 KD bulk RNA-sequencing.**

A, RT-qPCR analysis of *LSD1*, *ATOH1*, *NEUROG3*, and *HES1* RNA expression levels in HT29 untreated cells, or in cells treated with DMSO or DBZ. Significance determined by two-way ANOVA with Dunnett's multiple comparisons testing (****, P<0.0001). B, Barplot shows Log2FC in gene expression between LSD1 KD and EV in RNA-seq datasets from HT29 or SW480 cell lines. Orange bars denote significant changes in gene expression (FDR <= 0.05) and grey bars are not significant.

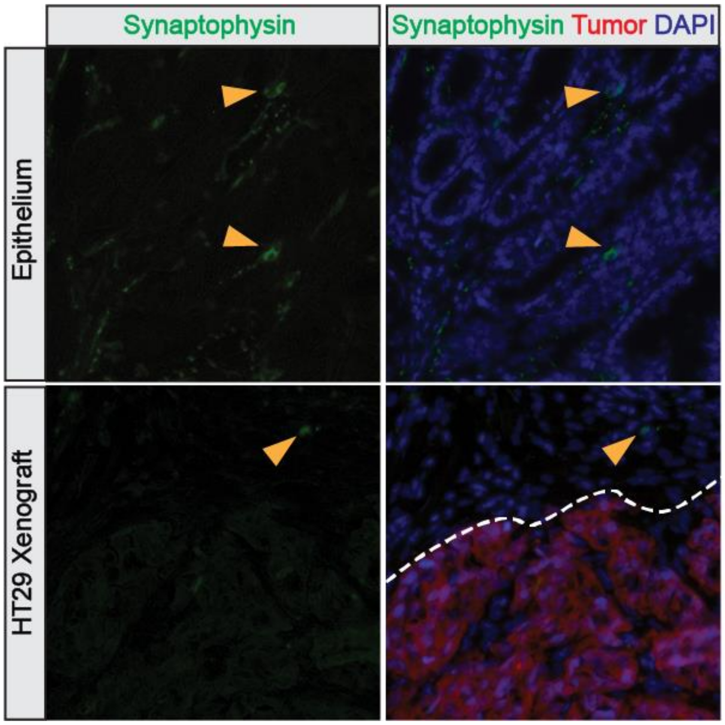


**Supplemental Figure 12: The HT29 cell line does not form Synaptophysin positive cells *in vivo*.**

Immunofluorescence images of primary xenograft tumors and adjacent normal colon. Images are representative of two biological replicates.

Barretina, J., Caponigro, G., Stransky, N., Venkatesan, K., Margolin, A.A., Kim, S., Wilson, C.J., Lehar, J., Kryukov, G.V., Sonkin, D.*, et al.* (2012). The Cancer Cell Line Encyclopedia enables predictive modelling of anticancer drug sensitivity. Nature *483*, 603-607.
Becht, E., McInnes, L., Healy, J., Dutertre, C.A., Kwok, I.W.H., Ng, L.G., Ginhoux, F., and Newell, E.W. (2018). Dimensionality reduction for visualizing single-cell data using UMAP. Nat Biotechnol.
Bergen, V., Lange, M., Peidli, S., Wolf, F.A., and Theis, F.J. (2020). Generalizing RNA velocity to transient cell states through dynamical modeling. Nat Biotechnol.
Butler, A., Hoffman, P., Smibert, P., Papalexi, E., and Satija, R. (2018). Integrating single-cell transcriptomic data across different conditions, technologies, and species. Nat Biotechnol *36*, 411-420.
Hafemeister, C., and Satija, R. (2019). Normalization and variance stabilization of single-cell RNA-seq data using regularized negative binomial regression. Genome Biol *20*, 296.
Hahne, F., and Ivanek, R. (2016). Visualizing Genomic Data Using Gviz and Bioconductor. Methods Mol Biol *1418*, 335-351.
Han, X., Zhou, Z., Fei, L., Sun, H., Wang, R., Chen, Y., Chen, H., Wang, J., Tang, H., Ge, W.*, et al.* (2020). Construction of a human cell landscape at single-cell level. Nature *581*, 303-309.
La Manno, G., Soldatov, R., Zeisel, A., Braun, E., Hochgerner, H., Petukhov, V., Lidschreiber, K., Kastriti, M.E., Lonnerberg, P., Furlan, A.*, et al.* (2018). RNA velocity of single cells. Nature *560*, 494-498.
Larsson, J. (2020). eulerr: Area-Proportional Euler and Venn Diagrams with Ellipses.
McCarthy, D.J., Chen, Y., and Smyth, G.K. (2012). Differential expression analysis of multifactor RNA-Seq experiments with respect to biological variation. Nucleic Acids Res *40*, 4288-4297.
McInnes, L., Healy, J., and Melville, J. (2020). UMAP: Uniform Manifold Approximation and Projection for Dimension Reduction.
Phipson, B., and Smyth, G.K. (2010). Permutation P-values should never be zero: calculating exact P-values when permutations are randomly drawn. Stat Appl Genet Mol Biol *9*, Article39.
Stuart, T., Butler, A., Hoffman, P., Hafemeister, C., Papalexi, E., Mauck, W.M., 3rd, Hao, Y., Stoeckius, M., Smibert, P., and Satija, R. (2019). Comprehensive Integration of Single-Cell Data. Cell *177*, 1888-1902 e1821.
Traag, V.A., Waltman, L., and van Eck, N.J. (2019). From Louvain to Leiden: guaranteeing well-connected communities. Sci Rep *9*, 5233.
Wang, Y., Song, W., Wang, J., Wang, T., Xiong, X., Qi, Z., Fu, W., Yang, X., and Chen, Y.G. (2020). Single-cell transcriptome analysis reveals differential nutrient absorption functions in human intestine. J Exp Med *217*.
Wolf, F.A., Hamey, F.K., Plass, M., Solana, J., Dahlin, J.S., Gottgens, B., Rajewsky, N., Simon, L., and Theis, F.J. (2019). PAGA: graph abstraction reconciles clustering with trajectory inference through a topology preserving map of single cells. Genome Biol *20*, 59.
